# Understanding the paradoxical mechanical response of in-phase A-tracts at different force regimes

**DOI:** 10.1101/854968

**Authors:** Alberto Marin-Gonzalez, Cesar L. Pastrana, Rebeca Bocanegra, Alejandro Martín-González, J.G. Vilhena, Rubén Pérez, Borja Ibarra, Clara Aicart-Ramos, Fernando Moreno-Herrero

## Abstract

A-tracts are A:T rich DNA sequences that exhibit unique structural and mechanical properties associated with several functions *in vivo*. The crystallographic structure of A-tracts has been well characterized. However, their response to forces remains unknown and the variability of their flexibility reported for different length scales has precluded a comprehensive description of the mechanical properties of these molecules. Here, we rationalize the mechanical properties of A-tracts across multiple length scales using a combination of single-molecule experiments and theoretical polymer models applied to DNA sequences present in the *C. elegans* genome. Atomic Force Microscopy imaging shows that phased A-tracts induce long-range (∼200 nm) bending. Moreover, the enhanced bending originates from an intrinsically bent structure rather than as a consequence of larger flexibility. In support of this, our data were well described with a theoretical model based on the worm-like chain model that includes intrinsic bending. Magnetic tweezers experiments confirm that the observed bent is intrinsic to the sequence and does not rely on particular ionic conditions. Using optical tweezers, we assess the local rigidity of A-tracts at high forces and unravel an unusually stiff character of these sequences, as quantified by their large stretch modulus. Our work rationalizes the complex multiscale flexibility of A-tracts, shedding light on the cryptic character of these sequences.

## INTRODUCTION

A-tracts are DNA sequences consisting of four or more consecutive A:T base pairs without the TA step. They are widespread across the genomes of both prokaryotic and eukaryotic organisms, including humans ^1–4^. Notably, the distribution of A-tracts along their genomes has proven to be essential for the proper organization of their genetic material with implications in transcription regulation ^5–9^. In addition, A-tracts have been shown to play an important role in recombination ^10, 11^, replication ^12^, antiviral response ^13, 14^ and stochastic gene silencing ^15^.

Remarkably, many of the functions of A-tracts have been linked to their particular structure and mechanical properties ^5, 10, 11, 16^. Regarding the structure, A-tracts are known to introduce a directional bend in the DNA helical axis ^11^. When two or more A-tracts are located in phase with the helical pitch, their bending adds constructively leading to a significant global curvature of the molecule ^17^. This curvature contrasts with the anomalously straight conformation reported for a single A-tract ^18, 19^. The most widely accepted solution to this conflict is the junction model, in which the bending is primarily localized at the edges of the A-tracts ^17^. Nevertheless, the precise bending mechanism in A-tracts remains a matter of debate^11, 20, 21^.

The reported mechanical properties of A-tracts are to some extent paradoxical. Early crystallographic studies suggested that A-tracts are stiff ^19, 22, 23^. This rigidity is in line with the high stretch modulus reported by recent molecular dynamics simulations of short (∼15 bp) duplexes containing an A-tract ^24, 25^, and can be attributed to the distinct structure of these sequences ^19, 24^. On the other hand, cyclization studies performed on duplexes longer than ∼100 bp revealed that A-tracts are not particularly rigid ^26, 27^. Other studies revealed that they are even more flexible than standard DNA sequences ^28^. This was supported by single-molecule experiments showing an enhanced curvature and looping probability in A-tracts ^29–31^.

Taken together, these results revealed a remarkable length-dependence of the mechanical properties of A-tracts and call for a unified comprehensive study. Such description should rationalize why these molecules appear rigid at scales of one helical turn, but flexible at scales longer than ∼100 bp. Moreover, a full characterization of the mechanical properties A-tracts should quantitatively distinguish their entropic bendability from their intrinsic static bending. This task is non-trivial in either structural or cyclization studies and relies on precise knowledge of the molecules’ trajectories over distances of hundreds of base pairs.

In this work, we study the mechanical properties of phased A-tracts from the genome of *C. elegans* (also referred to as hyperperiodic DNA) at multiple forces and length scales, at the single-molecule level using atomic force microscopy (AFM), magnetic tweezers (MT) and optical tweezers (OT). AFM imaging showed that phased A-tracts induce long-range bending on DNA molecules. The bending could be explained due to the presence of an intrinsically bent structure. Interestingly, MT experiments showed that at forces F<10 pN, phased A-tracts significantly soften the mechanical response of DNA molecules under low stretching forces. Whereas, OT experiments showed that at forces F>10 pN, the situation is reversed and the A-tracts confer DNA an unprecedented rigidity. Altogether, our work unveils the complex interplay between structural and mechanical properties of A-tracts across multiple force scales.

## RESULTS

### In-phase A-tracts induce intrinsic bending over long distances

In order to determine the mechanical properties of A-tracts in the absence of force, we used AFM to image DNA fragments of the genome of *C. elegans* containing in-phase A-tracts (Fig. 1, and **Experimental Section**). As a control, we imaged DNA molecules with heterogeneous sequence and similar length. AFM has proven to be an invaluable tool for characterizing DNA mechanical properties, providing a rigorous experimental test of DNA polymer models ^29, 32–34^. AFM imaging captured two-dimensional equilibrium conformations of the control (Fig. 2a) and phased A-tracts DNA molecules (Fig. 2b). As a first quantitative characterization, we computed the mean contour length of the control and the A-tracts molecules, obtaining values of 897±17nm (n=25, error is std) and 892±14nm (n=23), respectively. These values yield in both cases a helical rise of 0.34 nm bp^-1^, in agreement with the standard value of B-DNA ^35^.

**Figure 1.**
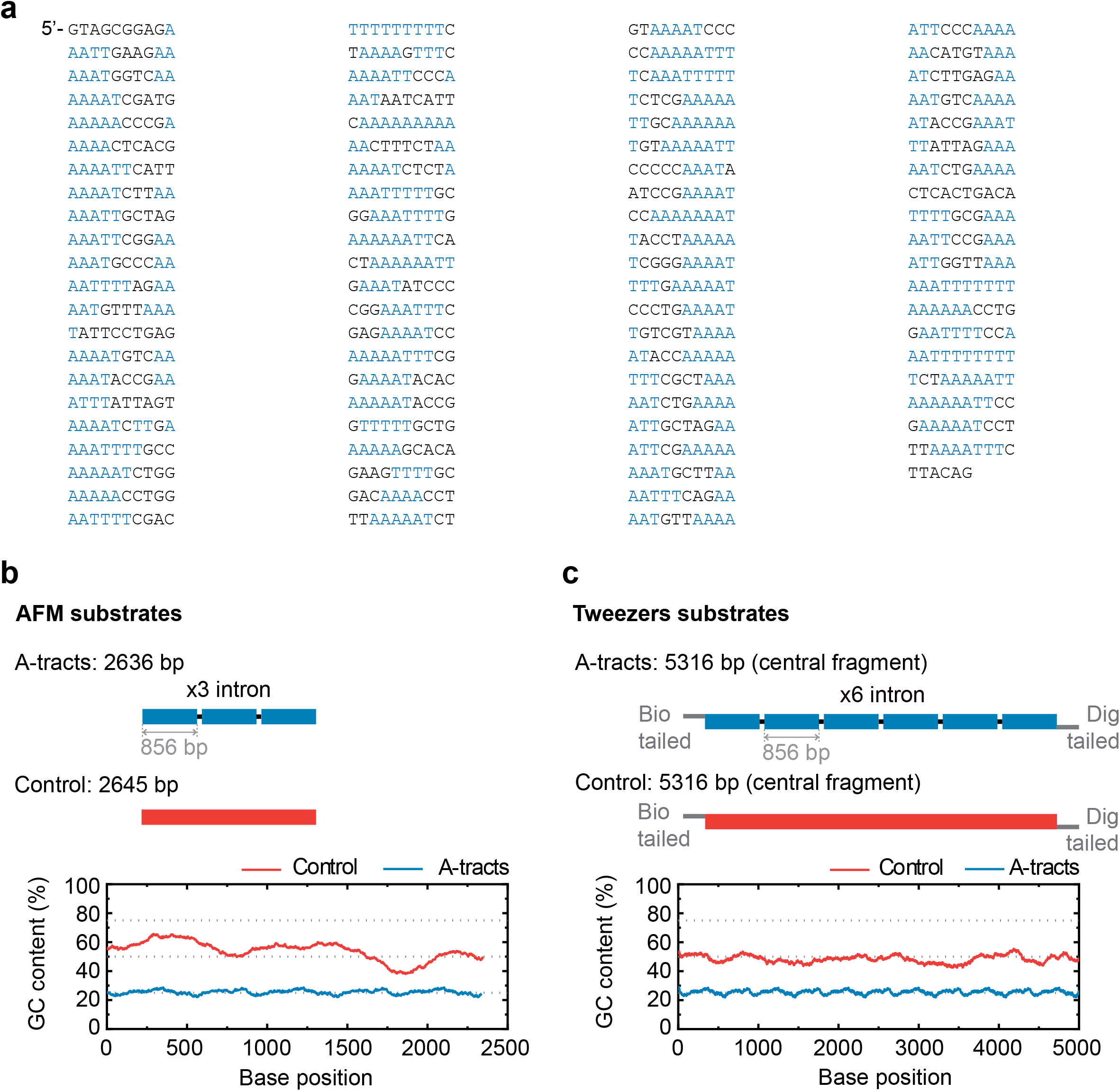
Sequence and overall organization of DNA molecules under study. (**a**) Phased A-tract sequence (intron) studied in this work as reported in ^30^ and Methods section. A-tracts (regions of four or more consecutive A’s and T’s without the TA step) are marked in blue. The sequence was written from the 5’ to the 3’-end in columns of 10 letters to highlight the ∼10 bp periodicity. (**b**) Top, the A-tract substrate for AFM imaging consisted on 3 repetitions of the intron, depicted by small blue blocks. A dsDNA molecule of heterogeneous sequence and similar number of bp was used as a control (red). Bottom, GC-contents of A-tract and control substrates was computed using a homebuilt software and selecting a running window of 300 bp. (**c**) Top, the DNA substrates for MT and OT contained six repetitions of the intron and were flanked by two oligonucleotides labelled with biotin and digoxigenin. A dsDNA molecule of heterogeneous sequence and similar number of bp was used as a control (red). Bottom, GC-content of A-tract and control molecules for tweezers experiments.

**Figure 2.**
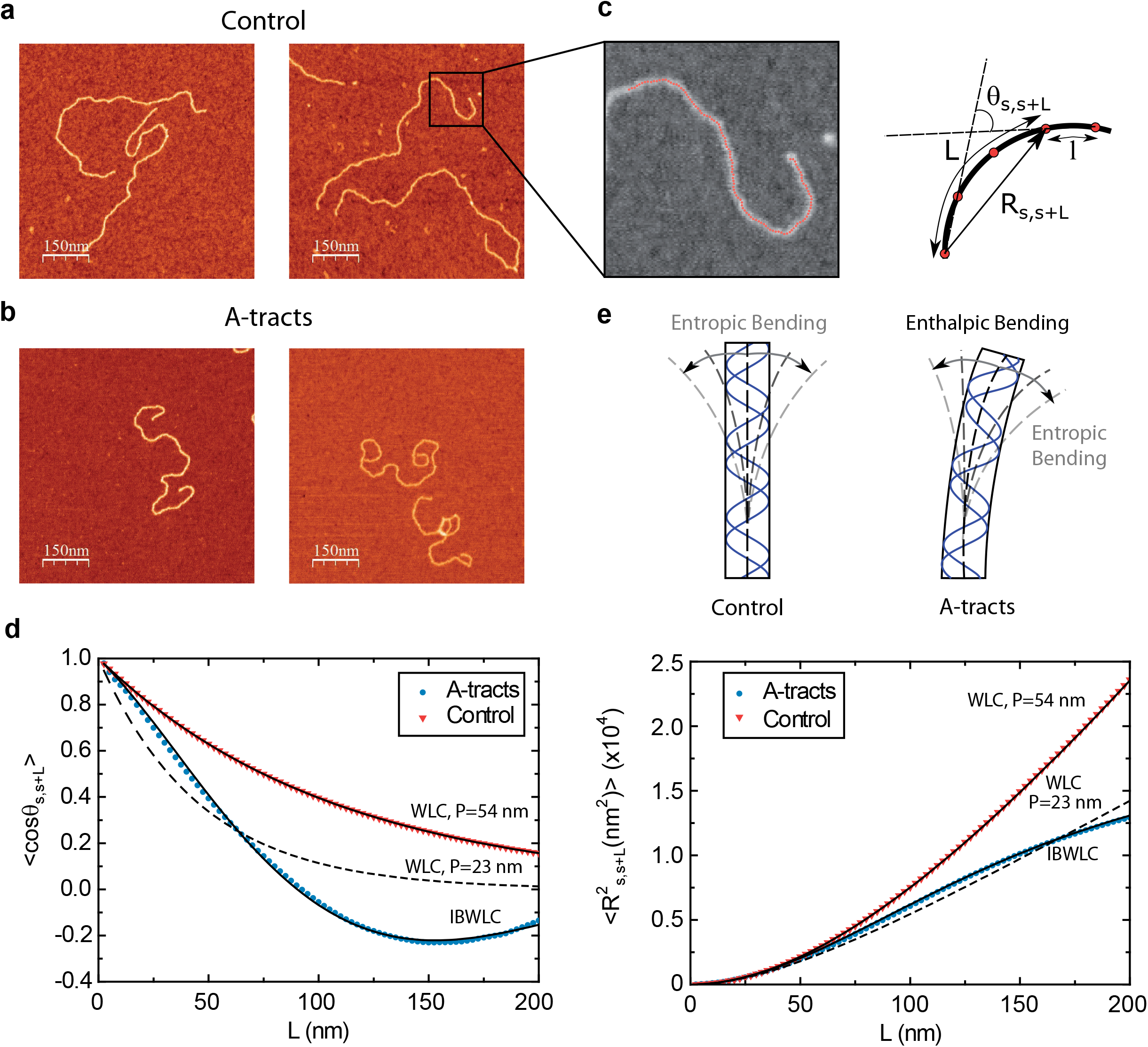
Mechanical properties of A-tracts at zero force. Representative AFM images of (**a**) control and (**b**) A-tract DNA molecules. (**c**) Detailed view of an individual DNA molecule deposited on mica surface. A series of dots (in red) separated by 2.5 nm were used to define the contour of the DNA molecules. Right, schematic representation of the measured quantities showing the angles between the tangents, *θ*_*s*,*s*+*L*_, and the distance, *R*_*s*,*s*+*L*_, for two points separated by a contour distance of *l*=7.5nm. (**d**) The quantities *cosθ*_*s*,*s*+*L*_ and 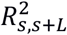, were averaged over all the traces, < *cosθ*_*s*,*s*+*L*_ > and 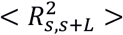, and plotted as a function of the contour distance separation between points, L. < *cosθ*_*s*,*s*+*L*_ > values of control and A-tract molecules were fitted to the WLC and IBWLC using **Eq. 2** and **Eq. 5**, respectively. This yielded a value of P = 54 ± 1 nm for control DNA and P =55 ± 1 nm, a = 17.4 ± 0.1 µm^-1^ for the A-tract molecule. Solid lines in the 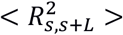 plot are not fits but the representation of **Eq. 3** and **Eq. 6** (WLC and IBWLC, respectively) using the fitting parameters obtained from the < *cosθ*_*s*,*s*+*L*_ > plot. Dashed lines represents best fits of the A-tracts data to the WLC model with P = 23 ± 1 nm. (**e**) Schematic representation of the WLC and IBWLC models. In the WLC model, the minimum energy corresponds to a straight conformation (zero enthalpic bending; dashed black line) but the molecule can be entropically bent (dashed grey lines) as a result of its bending flexibility. In the IBWLC model, the molecule is both enthalpically (curved dashed black line) and entropically (grey dashed line) bent.

A-tracts molecules appeared more bent than the control ones, in line with the findings reported in ^30^. In order to quantify the flexibility of these molecules, we firstly analysed their traces following a previously published protocol ^30, 34^. We calculated pairs of coordinates separated by a distance *l* = 2.5 nm that describe the contour followed by the adsorbed DNA molecules (Fig. 2c). From these traces, we obtained the angle, *θ*_*s*,*s*+*L*_, defined by the tangents at two points separated by a contour distance *L*; and the distance, 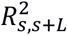, between these points (Fig. 2c). We then averaged 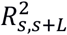 and *cosθ*_*s*,*s*+*L*_ over all the points of the trace and over hundreds of traces. Representing these two quantities, 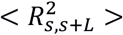 and < *cosθ*_*s*,*s*+*L*_ >, as a function of *L* allows direct comparison of AFM data with polymer models, see below(Fig. 2d).

The mechanical properties of DNA molecules are usually analysed in the context of the worm-like-chain (WLC) model, which describes polymers with harmonic bending energy

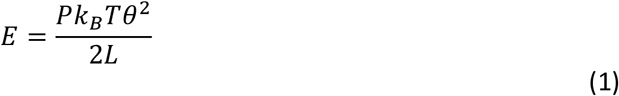

Where θ is the bending angle; *L* the contour separation as defined for the AFM traces; and *P* is the persistence length, which is proportional to the bending stiffness: *P* = *B*⁄*k*_*B*_ *T* ^32, 36^. Notice that the minimum energy configuration corresponds to θ = 0, that is, to straight molecules. Therefore, according to the WLC model, DNA molecules are bent solely by thermal fluctuations, i.e., they are *entropically bent*. The AFM data can then be fitted to the equations of the WLC in two dimensions:

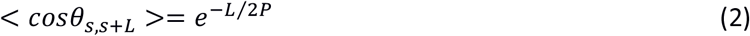

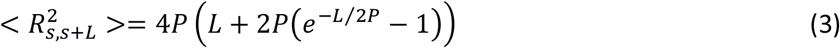

In agreement with previous works, our control data nicely fit to **Eq. 2** and **Eq. 3** with *P*= 54 ± 1 nm ^32, 34^ (Fig. 2d). However, A-tracts data deviated from the WLC behavior, and the best fit to **Eq. 3** provided an extremely low persistence length of *P*= 23 ± 1 nm (Fig. 2d, left panel). This deviation is even more remarkable for the cosine’s correlation, for which the WLC predicts an exponential decay (**Eq. 2**). In contrast, the cosine’s correlation of the A-tracts molecules reached negative values, with a minimum of ∼-0.2 around *L*= 150 nm (Fig. 2d, right panel). This demonstrated that, at zero force, the simple picture of the WLC model that assumes purely *entropically bent* polymers is not sufficient to describe the flexibility of A-tracts DNA molecules.

### An intrinsically-bent worm-like-chain model captures the long-range bending induced by in-phase A-tracts

We solved this discrepancy by developing a polymer model that incorporates intrinsic curvature into the WLC, as recently done for collagen fibers ^37^. This intrinsically-bent WLC (IBWLC) model has a simple, analytical solution that can be easily fitted to the AFM data, allowing to quantitatively decouple the bending contribution arising from thermal fluctuations (entropic contribution) from that coming from the intrinsic structure of the A-tracts (enthalpic contribution). This model is based on the assumption that the minimum of bending energy corresponds to a bent trajectory described by the arc of a circle of radius *R*_0_. This assumption has been widely used in the literature to explain that A-tracts induce a bend in the structure of the DNA ^11^, and cyclization ^21, 38^ and AFM data ^30^. This assumption is sketched in Fig. 2e and can be expressed mathematically as

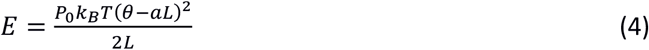

where *a* = 1/*R*_0_ is the intrinsic curvature, and *P*_0_ will be termed as *natural persistence length* of the molecule, which quantifies its resistance to bending similar to the WLC case. Notice that the WLC expression (**Eq. 1**) is recovered in a straightforward manner by making *a* = 0, that is, by deleting the intrinsic curvature; and by substituting *P*_0_ by *P*. From **Eq. (4)** we obtained the relevant < *cosθ*_*s*,*s*+*L*_ > and 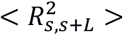 expressions for fitting the experimental data to the IBWLC, in a similar fashion to the WLC case ^32^:

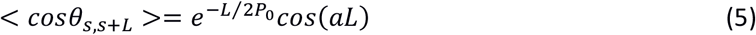

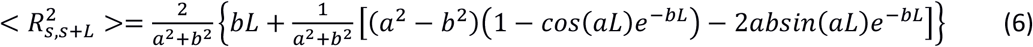

where we have defined *b* ≡ 1⁄(2*P*_0_) for convenience. A full derivation of the model can be found in ^37^ and in **Supplemental Material**.

A fit of the IBWLC model (**Eq. 5**) to the A-tracts data provided the fitting parameters *a* = 17.4±0.1 μm^-1^ and *P*_0_ = 55±1 nm (Fig. 2d, right panel). Using these values, we plotted the theoretical expression of 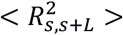 (**Eq. 6**) and found an excellent agreement with the experimental data (Fig. 2d, left panel). Remarkably, this agreement held for contour distances that extended up to at least 200 nm.

The IBWLC model enabled us to quantitatively dissociate the entropic and enthalpic contributions to the A-tracts flexibility. The value that we found for the intrinsic curvature, (*a* = 17.4±0.1 μm^-1^) was close to a previous estimation obtained of ∼10 μm^-1^ from AFM images^30^; and the persistence length essentially coincided with the control and with indirect measurements from cyclization studies on A-tracts ^26, 27^. Altogether, our results indicate that the A-tracts molecule has no enhanced bending flexibility. In other words, deviations from the minimum energy are energetically as costly as for the control molecule, as quantified by a similar persistence length for both molecules. On the contrary, the apparent larger bendability of this molecule stems from a purely *structural* or *static* deformation, its intrinsic bending.

### The low-force response of in-phase A-tracts molecules deviates from purely entropic models

AFM experiments show that the intrinsic A-tracts’ bending is responsible for the anomalous mechanical properties of these molecules at zero force. However, inside the cell the DNA is often subjected to mechanical stress, and to which extent the intrinsic bending of DNA affects its extension under mechanical force is unclear. We employed MT to explore the mechanical response of phased A-tracts sequences below 6 pN. To this end, we took force-extension curves of DNA molecules containing 6 repetitions of an A-tract sequence found in *C. elegans* (Fig. 1c) and compared them with those of a DNA control with heterogeneous sequence and similar number of base pairs (100 mM NaCl, see **Experimental Section** for details). Averaged force-extension curves showed that below 5 pN, A-tracts molecules presented significant lower extensions than those of the control (Fig. 3a). Importantly, this decrease in end-to-end distance of the molecule was also found at zero force in the AFM experiments and was rationalized to be an effect of the bends (**Eq. 3** and Fig. 2d, left panel). Notably, this effect virtually disappears at a force of ∼5 pN, where both molecules show a similar extension and practically reach their full contour length (∼1.8 μm).

**Figure 3.**
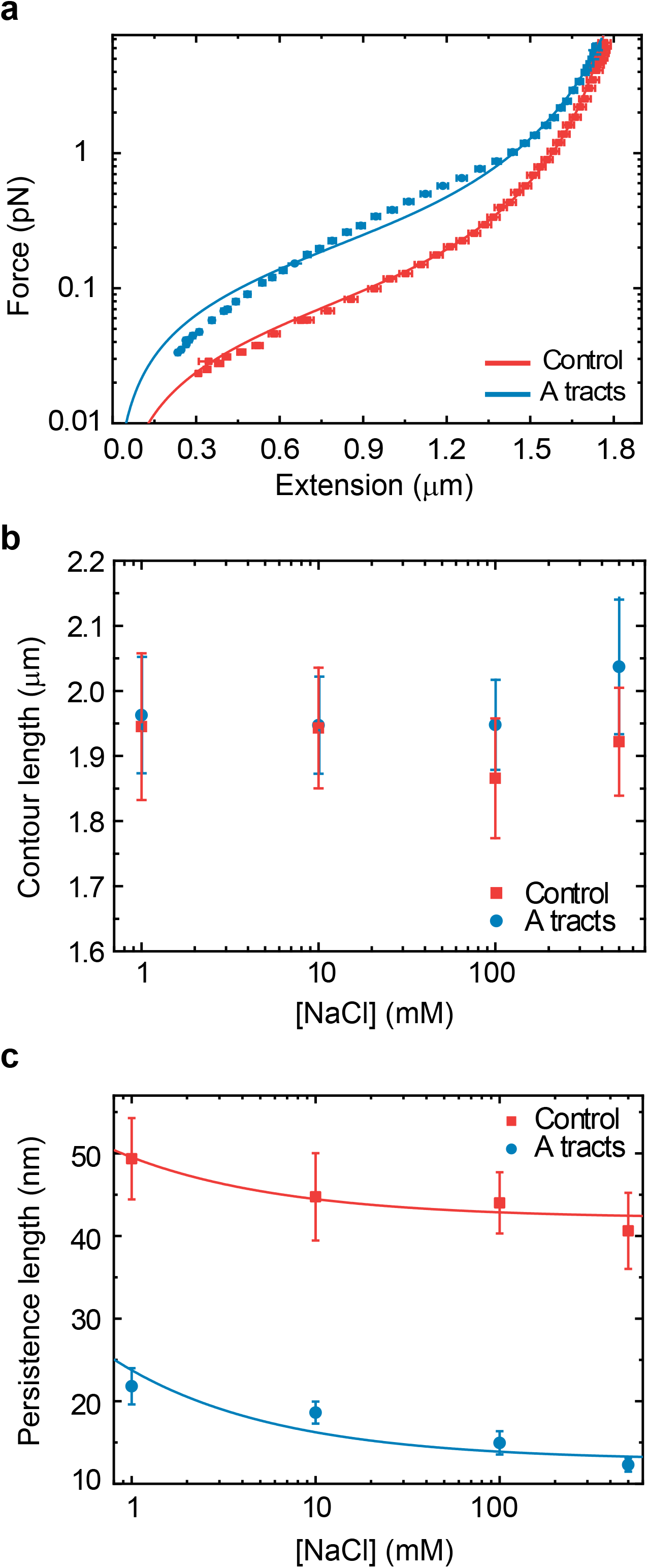
Mechanical response of A-tracts to forces below 10 pN. (**a**) Average force-extension curves of A-tracts and control DNA molecules obtained with magnetic tweezers (100 mM NaCl, mean ± sem of N>20 molecules). Solid lines are fits to the WLC formula accounting for low force corrections. (**b**) Contour length values (mean ± std) obtained from the fits to the force-extension curves of control (red) and A-tracts (blue) molecules with **Eq. 7** at different salt concentrations. (**c**) Relation between the persistence length *P* and monovalent salt concentration (mean ± sd). The lower errors for A-tracts are the result of the larger difference between force-extension curves at low persistence lengths and at low forces. The data was fit to the Barrat and Joanny model, *P* = *P*_∞_ + *mc*^−1/2^, where *P*_∞_ is the persistence length in the limit of high salt concentration, *m* is a fitting parameter, and *c* is the salt concentration ^68^. We found *P*_∞_ = 42 ± 1 nm, *m* = 7 ± 2 nm·mM^1/2^ for control molecules and *P*_∞_ = 14 ± 1 nm, *m* = 11 ± 4 nm·mM^1/2^ for A-tracts, indicating that A-tract intrinsic curvature is not strongly dependent on monovalent cations concentration.

In order to quantify the mechanical properties of A-tracts we resorted to the WLC interpolation formula ^39^:

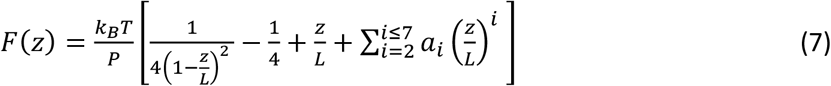

where *z* is the extension of the molecule, *F* the applied force, *P* the persistence length and the phenomenological constants *a*_*i*_ are a *a*_2_ = −0.5164228, *a*_3_ = −2.737418, *a*_4_ = 16.07497, *a*_5_ = −38.87607, *a*_6_ = 39.49944 and *a*_7_ = −14.17718.

As expected, force extension curves of control DNA were well described by the WLC model (Fig. 3a, red symbols) and yielded a persistence length of *P* = 44 ± 4 nm, in agreement with previous measurements ^34, 40^. However, the WLC model could not properly fit the A-tracts data over the entire force range (Fig. 3a, blue symbols). At forces below 1pN the data were best described by a *P*= 15 ± 1 nm, whereas in the 1-4 pN force range data were better fitted to a higher persistence length of *P*=21±8 nm. The deviation of the A-tracts data from the WLC model likely reflects the presence of intrinsic bends in the DNA, given that the model describes polymers that are purely entropically bent. Consequently, we propose that, similar to the zero force case (Fig. 2d), the deviation from the WLC together with the anomalously small persistence length constitute a signature of the presence of intrinsic bends. Therefore, in absence of a better model, we will use here the WLC and quantify the bent character of the A-tracts molecules from the low value of their persistence length. Future theoretical works might provide a more precise measurement of the A-tracts intrinsic bending from our MT experimental data by employing.

It has been argued that A-tracts bending may be the consequence of the interaction of monovalent ions with the minor groove ^41, 42^. Conversely, other works favour an intrinsic ion-independent bending model as a result of the distinct base-pairing and stacking of AT-rich domains ^43, 44^ and even suggest a loss of curvature with the increase of monovalent salt concentration ^45^. In order to elucidate the role of monovalent ions on A-tracts bending, we obtained additional force-extension curves at NaCl concentrations of 1, 10 and 500 mM (**Fig. S2**) and compare them with control sequences. Fits from the WLC model resulted in similar values of contour lengths of both molecules and for all salt concentrations (Fig 3b). In addition, we found that the persistence length of the A-tracts and the control molecules followed a similar decaying trend with increasing NaCl concentrations, as it has been reported for DNA molecules of arbitrary sequence ^34, 46^ (see **Supplementary Information**, and Fig. 3c). However, in all cases, the WLC could not fit the force-extension data of A-tracts sequences (**Fig S2**), as shown for the 100 mM NaCl case (Fig. 3a). Therefore, we conclude that A-tracts bending is intrinsic to the structure of the molecule and does not rely on monovalent ions.

### A-tracts present a high stretching rigidity

Further to our analysis in the entropic low force regime, we then asked about the mechanical response of A-tracts in the enthalpic regime by determining their stretching rigidity. We used optical tweezers (OT) to obtain force-extension curves of the control and the A-tract molecules in a buffer containing 100 mM NaCl (see **Experimental Section** for further details). As expected for random DNA sequences, we observed a smooth overstretching transition at ∼60 pN for the control molecule ^47–49^ (**Fig. S3**). However, the overstretching transition of the A-tract molecule started at a lower force, showing characteristic periodic saw-tooth pattern with six repetitions. These repetitions likely correspond to the six copies of the intron (Fig. 1c), in line with previous OT experiments reporting reproducible sequence-dependent unpeeling of overstretched DNA^50^.

To quantify the stretching response of A-tracts molecules, we first fitted force-extension data in the 1-10 pN range to the inextensible WLC model (Fig. 4a**, inset**) ^34^:

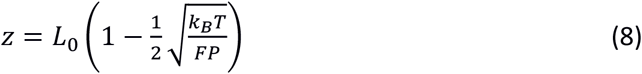

**Figure 4.**
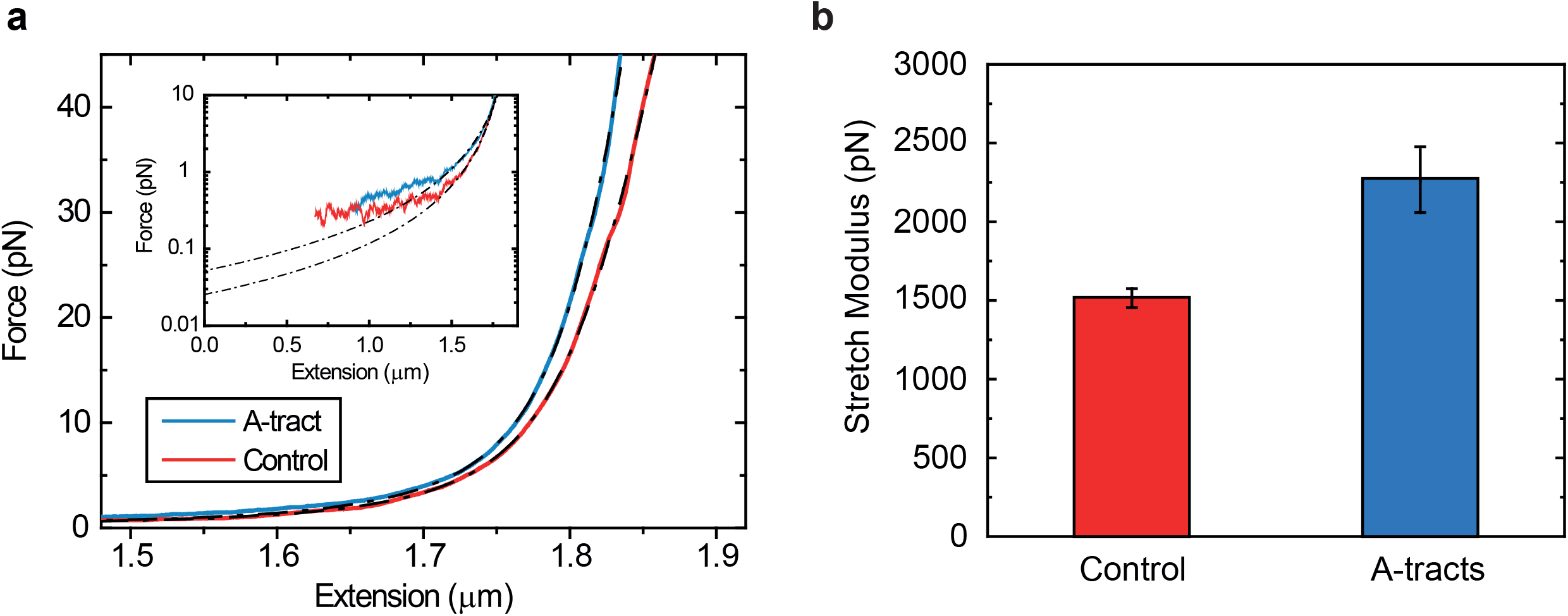
Mechanical response of A-tracts to forces above 10 pN. (**a**) Representative force-extension curves of control and A-tract molecules measured with OT. Dashed lines represent the fits to the eWLC of the control and A-tract molecules in the 10-45 pN range. The values of these particular fits were *L*_0_ = 1852 nm, *P* = 44 nm, *S* = 1704 and *L*_0_ = 1851 nm, *P* = 39 nm,*S* = 2802 nm for the control and A-tract molecules respectively. Note that the eWLC does not fit well the A-tracts data in the 1-10 pN interval. Inset, the same experimental curves are represented using a logarithmic force scale and showing the fits to the WLC in the 1-10 pN range. The values of these fits were *L*_0_ =1863 nm, *P* =41 nm and *L*_0_ =1916 nm, *P* =20 nm for the control and A-tracts data respectively. (**b**) Average stretch modulus (S) of control and A-tracts molecules (see Table 1) at 100 mM NaCl. Error bars represent the standard error of the mean of the fits.

**Table 1.** Mechanical parameters of A-tracts and control molecules from MT and OT experiments. The parameters shown are the average of the fits and the errors are the standard deviation of the mean. *L*_0_ and *P*_*WLC*_ were obtained by fitting the MT (OT) data to Eq. 7 (Eq. 8) in the *F* < 1 pN (1 < *F* < 10 pN) range. *P*_*eWLC*_ and *S* were calculated by fitting the OT data to Eq. 9 in the 10-45 pN range.

| Magnetic Tweezers |  |  |  |  |  |
| --- | --- | --- | --- | --- | --- |
| [NaCl] (mM) | Molecule | $L_0$ (nm) | $P_{WLC}$ (nm) | $P_{eWLC}$ (nm) | $S$ (pN) |
| 1 | A-tract | $1936 \pm 30$ | $22 \pm 1$ | | |
| | Control | $2013 \pm 35$ | $49 \pm 1$ | | |
| 10 | A-tract | $1928 \pm 20$ | $19 \pm 1$ | | |
| | Control | $1964 \pm 40$ | $45 \pm 1$ | | |
| 100 | A-tract | $1913 \pm 10$ | $15 \pm 1$ | | |
| | Control | $1846 \pm 25$ | $44 \pm 1$ | | |
| 500 | A-tract | $1930 \pm 20$ | $12 \pm 1$ | | |
| | Control | $1970 \pm 20$ | $41 \pm 1$ | | |
| Optical Tweezers |  |  |  |  |  |
| [NaCl] (mM) | Molecule | $L_0$ (nm) | $P_{WLC}$ (nm) | $P_{eWLC}$ (nm) | $S$ (pN) |
| 50 | A-tract | $1891 \pm 5$ | $19 \pm 1$ | $38 \pm 3$ | $2410 \pm 160$ |
| | Control | $1832 \pm 4$ | $44 \pm 2$ | $51 \pm 4$ | $1660 \pm 110$ |
| 100 | A-tract | $1870 \pm 9$ | $19 \pm 1$ | $43 \pm 3$ | $2270 \pm 170$ |
| | Control | $1836 \pm 2$ | $42 \pm 1$ | $46 \pm 2$ | $1520 \pm 70$ |
| 500 | A-tract | $1841 \pm 3$ | $22 \pm 1$ | $38 \pm 3$ | $2370 \pm 170$ |
| | Control | $1829 \pm 2$ | $42 \pm 1$ | $46 \pm 3$ | $1580 \pm 70$ |

The values obtained for the control and the A-tracts molecule were *L*_0_= 1836 ± 2 nm, *P*= 42 ± 2 nm and *L*_0_= 1870 ± 9 nm, *P*= 19 ±1 nm, respectively, in agreement with the AFM and MT results (Fig. 2d **and Table 1**).

With the aim of characterizing the A-tracts mechanical properties at forces we then resorted to the extensible WLC (eWLC) model, which considers that the molecule can be elongated beyond its contour length ^51, 52^. According to the eWLC, the extension of a DNA molecule depends on the applied force as:

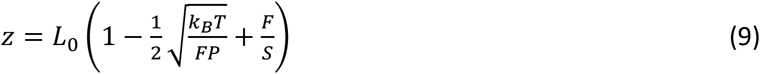

where *S* is the stretch modulus and quantifies the enthalpic elongation of the molecule. We used this expression to fit the force-extension curves of the control molecule in the 10-45 pN range. Remarkably, in this range of forces, **Eq. 9** fits well the experimental data (Fig. 4a). We obtained *L*_0_= 1824 ± 3 nm, *P* = 46 ± 2 nm and *S*= 1520 ± 70 pN for the control molecule, in agreement with the exhaustive literature on DNA flexibility ^34, 50, 53^ (**Table 1**). The values of the fitting parameters obtained for the A-tracts molecule were *L*_0_= 1805 ± 8 nm, *P*= 43 ± 3 nm, which is similar to that of the control molecule and *S*= 2270 ± 170 pN, which interestingly, is ∼50% higher than the enthalpy rigidity found for the control molecules (Fig. 4b). Consistently, similar results were obtained at NaCl concentrations of 50 mM and 500 mM (**Fig. S4** and **Table 1**).

## DISCUSSION

### A comprehensive picture of the mechanical properties of A-tracts

Our results can be brought together to build a comprehensive picture of the multiscale flexibility of A-tracts (Fig. 5). At zero force, the in-phase bends of the A-tracts result in a molecule with preferred curvature, as observed in the AFM images (Fig. 5a). These are responsible for the measurement of a distinctively small persistence length in these molecules, as found in MT and OT experiments at forces <5 pN. We propose that under an external force, these bends are gradually opened, allowing the molecule to eventually extend to its full contour length. Remarkably, such process of straightening the bends would represent an additional source of flexibility that is not captured by the WLC model, which only considers entropic elongation, resulting in a low value of the persistence length (Fig. 5b). This would explain the deviations from the WLC model observed in force-extension curves below forces of 10 pN (Fig. 3a). According to our data, at forces >10 pN), the bends characteristic of A-tracts would have been practically straightened and both the control and the A-tracts molecule show similar values of extension and persistence length (Fig. 3a, Fig. 4a, Fig. 5c and **Table 1**). Interestingly, this idea has been previously proposed in the context of the theoretical *kinkable WLC* model ^54^, in which a kinked elastic polymer under force would present a small effective persistence length at low forces and would recover its *natural persistence length* in the high force limit. To the best of our knowledge this is the first experimental demonstration of such puzzling effect.

**Figure 5.**
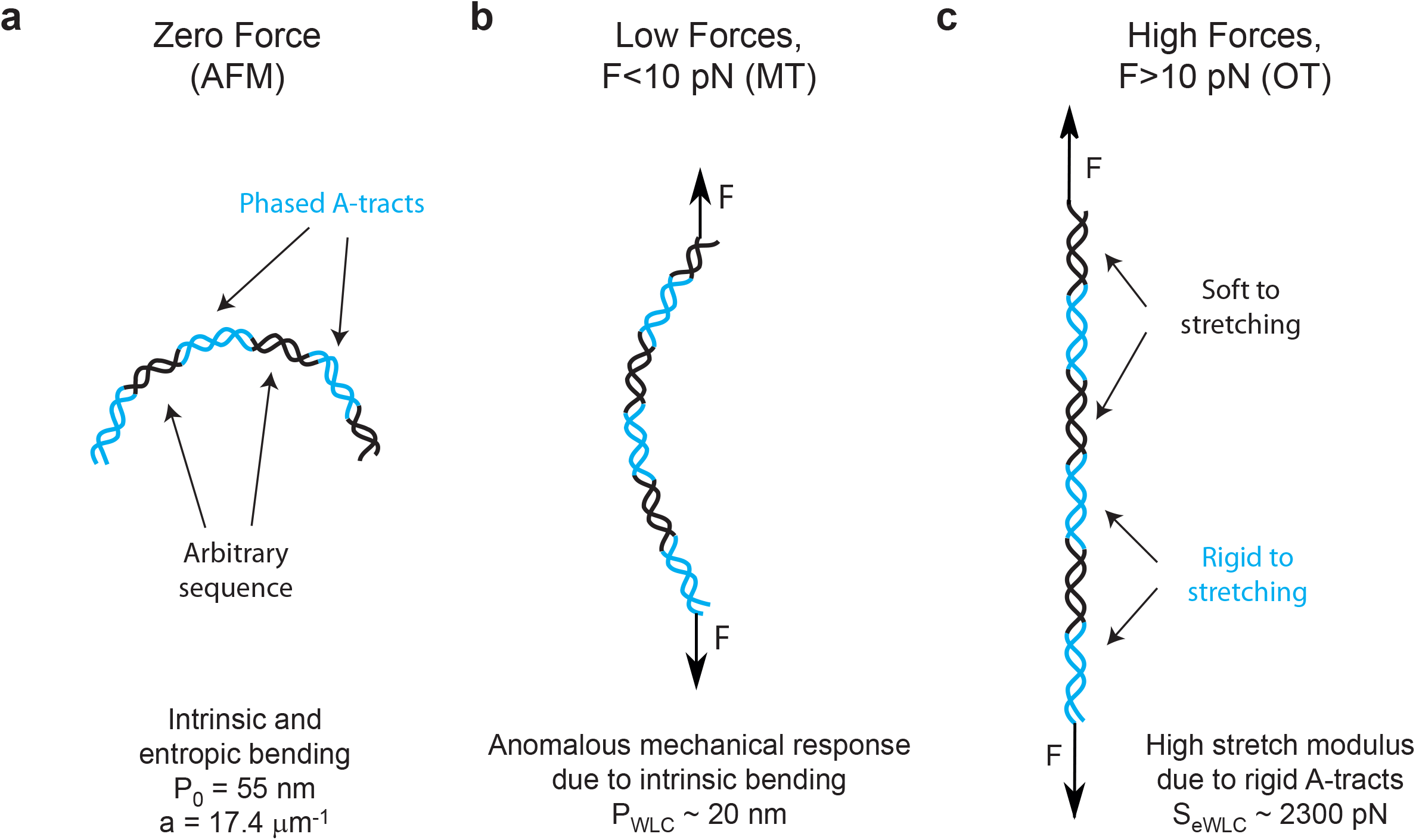
Schematic representation of the proposed model of the mechanical properties of A-tracts. **(a)** In the absence of stretching forces, molecules containing phased A-tracts (blue regions) appear largely bent as a consequence of the local structural bends. **(b)** Forces <10 pN align and straighten the bends along the pulling coordinate, which results in a significant deviation of the force-extension curves from the WLC model. **(c)** At forces >10 pN the bends are fully extended and the local elastic response of A-tracts, is exposed. This force regime unveiled the extraordinary enthalpic rigidity of A-tracts.

According to our data and model, at forces F>10 pN, the A-tracts molecules would be practically straightened and the local stretching rigidity of the A-tracts evaluated (Fig. 5c). We found an unusually high value for the stretch modulus of A-tracts of ∼2300 pN (Fig. 4b), in line with the one (∼2800 pN) reported for a ∼15 bp long poly-A sequence in a recent theoretical work ^24^ and in agreement with previous simulations that predict a high stretching rigidity for these sequences ^25, 55^. Moreover, this finding confirms the high rigidity usually attributed to these sequences in structural studies ^19, 22, 56^. A possible explanation for the high A-tract stiffness has been previously proposed in the context of the *crookedness model* for arbitrary DNA sequences ^24^. Inside the A-tract the base pair centres are almost perfectly aligned with the helical axis and thus, the molecule can only elongate by unstacking its base pairs, which is energetically very costly. Conversely, the large stretch modulus found here can be regarded as indicative of a particular structure of the A-tracts that differs from the canonical B-DNA, in the same way as the softer stretching response of dsRNA can be ascribed to its A-form ^57–59^.

### Biological Implications

A-tracts appear enriched in the genomes of diverse organisms ^1–4^ and their particular structures and mechanical properties may play a role on several biological processes including recombination, replication and DNA packaging ^10, 11^. In many of these processes, A-tracts are thought to operate by stabilizing the formation of DNA tertiary structures. For instance, A-tracts have been shown to localize supercoils ^60^ and might facilitate the formation of loops at regulatory regions ^11^. In addition, short phased A-tracts might stabilize the highly bent structure adopted by DNA in the nucleosome ^8, 61^ and likely contribute to DNA packaging in the bacterial nucleoid ^3^. However, somewhat surprising, in some other cases A-tracts seem to inhibit bent DNA conformations. Possibly, the most prominent example are poly-A stretches longer than 10 bp, which are thought to resist strong bending and have been shown to significantly destabilize nucleosomes ^5, 6^.

Based on our results, we may propose an explanation for this apparent contradiction on A-tracts flexibility, with views on achieving a better understanding of the biological function of these sequences. We have shown that phased A-tracts greatly bend the DNA; however the tracts themselves appear rigid at high forces. This finding suggests that the relative magnitude of these two effects - intrinsic bending and A-tract rigidity – could be modulated by tuning the length and distribution of the A-tracts. In-phase A-tracts shorter than 10 bp would amplify the intrinsic bending, whereas an individual A-tract longer than ∼10 bp would greatly stiffen the DNA. The former should then be involved in bent DNA conformations, such as in supercoils or nucleosomes; whereas the latter would be preferred for rigid sequences that preclude strong distortions in the DNA. This idea is in line with the findings reported in the literature ^5, 8, 60–62^.

Furthermore, the considerations above raise the question of which of the DNA mechanical properties are interrogated during nucleosome. In particular, if the mechanical properties of poly-A stretches are the responsible for their nucleosome depleting character, one would expect a high bending stiffness of these sequences. On the contrary, what we found is that A-tracts are stiff in terms of stretching, but not bending. In addition, a recent study on the flexibility of poly-A stretches does not support the generalization that these sequences are always resistant to bend ^31^. Therefore, we propose that stretching, which reflects a short-scale deformation ^24^, could play a more relevant role in nucleosome stabilization than the WLC bendability characteristic of long distances of up to several hundreds of nanometers. A similar idea has been proposed in simulation studies ^24, 25^. The high positive charge of the histones could well induce tight bends in the DNA. However, the precise adaptation of the DNA molecule to the shape imposed by the histones relies on very particular local motions of the DNA ^25, 63, 64^. A high stretch modulus might be indicative that these local motions are precluded. The hypothesis that bending is not determinant for nucleosome stabilization is supported by the structure of a nucleosome core particle containing a 16 bp-long poly-A. This tract is able to bend, but it adopts a distorted configuration that ultimately destabilizes the nucleosome ^65^. Interestingly, CpG islands, another well-known example of nucleosome destabilizing sequence ^66^, have also been reported to show a large stretch modulus of ∼1800 pN ^67^. Finally, in addition to its role in nucleosome positioning, A-tract rigidity would mechanically stabilize the structural features of these sequences, which constitute a recognition mechanism in DNA-protein interactions ^8, 16^.

## SUMMARY

In this work we have investigated the mechanical properties of A-tracts at the single-molecule level using atomic force microscopy (AFM), magnetic tweezers (MT) and optical tweezers (OT). Our AFM measurements reveal the intrinsic curvature of these sequences, as manifested by the deviation of their mechanical properties from the WLC model. We hence used a variation of this model that accounts for polymers with a preferred curvature. This model described our AFM experimental data with exceptional accuracy, and allowed us to quantify the intrinsic and entropic bending of these molecules. In addition, we employed MT and OT to characterize the response of A-tracts under the action of an external force. At low forces the mechanical response of A-tracts is described by a persistence length of ∼20 nm, a hallmark of bent DNA. This reduction of the persistence length of A-tracts was preserved for a wide range of NaCl concentrations, supporting that bending of A-tracts does not rely on the interaction with ions.

Interestingly, OT experiments revealed a ∼1.5-fold increase in the stretching modulus compared to random sequence DNAs. This exceptional stretching stiffness of A-tracts likely reflects the unusual local rigidity of these sequences which has been linked with their efficiency in nucleosome depletion. Our results reconcile contradictory views on A-tracts flexibility and stress the need of appropriate models and techniques to disentangle all the different mechanical deformations that may be involved across wide length scales at different mechanical forces. From a broader perspective, our comprehensive study on the mechanical properties of A-tracts might shed light on the intricate relation between sequence dependent flexibility and function in the double helix.

## ACKNOWLEDGEMENTS

The authors acknowledge the computer resources, technical expertise and assistance provided by the Red Española de Supercomputacion at the Minotauro Supercomputer (BSC, Barcelona). We thank Andrew Fire (Stanford University, USA) and Ralf Seidel (University of Leipzig, Germany) for providing us biological material required for the fabrication of the DNA molecules.

## EXPERIMENTAL SECTION

### Synthesis of DNA molecules for single-molecule experiments

In order to study the mechanical properties of A-tracts at the single-molecule level, we considered a hyperperiodic sequence of 856 bp from the *C. elegans* genome ^69^. This segment corresponds to the fourth intron of the gene F54C4.1 that encodes the *C. elegans* ortholog of human mitochondrial ribosomal protein L40. We will refer to this sequence as the *intron* (Figure 1a). The intron was PCR-amplified from the pPD167.57 plasmid, with the oligonucleotides 58.F Bam-Xho-Psi intron4 and 59.R Apa-Eco-Sal intron4 (**Table S1**). After digestion, the PCR product was electrophoresed on a 1% agarose gel, extracted (QIAGEN Gel Extraction Kit), and cloned into the pNLrep plasmid. This process was performed several times to obtain plasmids with 1 to 6 copies of the intron, and constitutes the basis to build the molecules needed for single molecule studies (Figure 1b, c).

### Synthesis of DNA molecules for AFM experiments

The A-tract substrate for AFM measurements was made of three copies of the intron, resulting in a 2636 bp molecule (A-tract AFM substrate, **Table S2**). This fragment was obtained by digestion of the respective plasmid with XhoI and EcoRV enzymes. As a control, a fragment of DNA of 2645 bp (control AFM substrate, **Table S2**) with a 54% content of GC (Figure 1b) was obtained by digestion of pNLrep-0BspQI site plasmid (previously cloned in the laboratory from pNLrep plasmid) with SalI and ScaI enzymes. Fragments coming from digestion or PCR were electrophoresed on a 1% agarose gel and extracted. DNAs were never exposed to intercalant dyes or UV radiation during their production and were stored at 4°C.

### Synthesis of DNA molecules for MT and OT experiments

We fabricated DNA molecules of 5316 bp containing 6 copies of the intron (A-tract tweezers substrate, **Table S2**). The central fragment of the molecule is flanked by oligonucleotides labelled either with digoxigenin (3’ end) or biotin (5’ end) that specifically bind either to a glass surface covered with Anti-digoxigenin (Roche) or to superparamagnetic beads (MyOne, Dynabeads) covered with streptavidin (Figure 1c). Labelled oligonucleotides were fabricated based on a previously published method ^70^ Briefly, 27P-XhoI-A and XbaI-A oligonucleotides (**Table S1**) were biotin or digoxigenin tailed using Terminal Transferase (NEB) and either BIO-dUTP or DIG-dUTP, respectively. The modified oligonucleotides were purified using a Qiaquick nucleotide removal kit (Qiagen) and hybridized respectively with 26XhoI-B or 88.XbaI C ApaI (**Table S1**). The central fragment was digested with XhoI and ApaI enzymes and ligated overnight to the two hybridized tailed oligonucleotides using T4 DNA ligase (NEB). Excess of oligonucleotides was removed using Microspin S-400 columns. As a control molecule (**Table S2**), we chose a fragment of DNA with a homogeneous GC content with the same length (Figure 1c). This fragment was PCR-amplified with the oligonucleotides 89.F lambda 40002 XhoI and 90.R lambda 45263 ApaI using Lambda DNA (NEB) as a template. This region was selected by running a homemade software that computes the GC content of a given sequence and selecting a running window of 300 bp. The PCR product was digested, electrophoresed on a 1 % agarose gel, extracted, and ligated with the same tailed oligonucleotides used to produce the A-tracts molecule.

A-tract molecules presented a low GC-content (∼20%) with periodic patterns that arise from the three and six repetitions of the intron present in the AFM and Tweezers’ substrates, respectively. In contrast, the GC-content of the control molecules are ∼50% (Figure. 1b, c). Importantly, all A-tracts constructs showed anomalous gel migration (Fig. S1), as expected for phased A-tracts ^30^.

### Atomic Force Microscopy Measurements

A 20 ml solution containing 0.3 nM of DNA in 100 mM NaCl, 10 mM Tris-HCl pH 8 and 15 mM MgCl_2_. was deposited on freshly-cleaved mica and incubated for ∼30 s. Then, the sample was washed with 4 mL of Milli-Q water and dried under nitrogen air-flow. Imaging conditions were similar to the ones described in a previous work ^34^. Images were taken with an AFM from Nanotec Electronica S.L. with PointProbePlus tips (PPP-NCH Nanosensors) and using tapping imaging mode in air. Sample images were obtained at a resolution of 1.46 nm perpixel and processed with WSxM freeware ^71^. Contour lengths were computed using the WSxM software and persistence lengths were obtained using the tracing routine described in ^30, 33^ by taking 290 nm traces with 2.5 nm point-to-point (*l*) separation.

### Magnetic Tweezers Measurements

We used a Magnetic Tweezers setup based on an inverted optical microscope illuminated by nearly monochromatic LED light to track micrometer-sized superparamegnetic beads tethered to the surface by the DNA molecule of interest ^72, 73^. The spatial coordinates of the beads are extracted by videomicroscopy analysis of two-dimensional correlation (xy coordinate) and by the analysis of the pattern of diffraction rings (z coordinate). Forces in the range of 0.03 pN to pN are applied by approximating two vertically-aligned magnets (Supermagnete, W-05-N50-G) separated by a gap of 1 mm. The force applied at a given magnet position was determined from the average extension of the molecule and from the analysis of the Brownian excursions in Fourier space. In addition, motion blur and the ensuing overrating of the force was minimized by tracking at high frequencies (400 Hz). Force-extension curves were obtained by sampling the average extension at a constant force. Molecular extensions were corrected by subtracting the extension at zero force. Double-tethered beads were discarded from our measurement attending to their characteristic rotations-extension response. In addition, we discarded DNA-beads showing large off-center attachment to prevent underrating the persistence length, an artifact previously reported ^74, 75^. Beads with large off-center attachment were identified from the projected circle in the xy plane when magnet turns are applied. All the experiments were performed in a buffer composed of 10 mM Tris-HCl pH 8.0, 1 mM EDTA and supplemented with NaCl at the quoted concentration.

### Optical Tweezers Measurements

Optical tweezers experiments were performed with a highly stable miniaturized dual-beam setup ^76^. Individual DNA constructs labelled with biotin at one end and digoxigenin at the other (see above) were attached between a streptavidin-covered bead held by suction on top of a micropipette and an anti-digoxigenin-covered bead located at the optical trap (force sensor). Axial forces were applied to single DNA molecules by displacing the pipette relative to the optical trap. Force-extension curves were obtained at 500 Hz of sampling frequency by moving the optical trap at a pulling rate of 200 nm s^-1^ with a spatial resolution of 1 nm and a force resolution of 1 pN. Our optical tweezers setup controls the distance between the trap center and the pipette (*X*_total_), which is different from the end-to-end distance of the DNA molecule (*X*_end–end_). The end-to-end distance was calculated as X_end−end_=X_Tot_−(F/k), where *F* is the force applied to the system, *k* the stiffness of the trap, and *F/k* corresponds to the distance (nm) moved by the bead out of the trap center (bead position). We assume a linear spring restoring force. The trap stiffness under our experimental conditions was *k* = 0.135 ± 0.0043 pN nm^−1^. Experiments were done in a buffer of 10 mM Tris-HCl pH 8.0, 1 mM EDTA with the concentration of NaCl specified in the text. Raw data was processed by computing a running average in windows of 100 points.

