## Supplementary Information for "Understanding the paradoxical mechanical response of in-phase A-tracts at different force regimes"

### Supporting Information

1. **Theory Model**
2. **Determining the role of monovalent ions on A-tract bending**
3. **Supporting Figures**
4. **Supporting Tables**

### Theory Model

#### 1. Worm-Like Chain (WLC) Model

It is helpful to start with the simpler case of the WLC model. Here we will follow the notation and the derivation from refs [1, 2]. According to the WLC model, the bending energy of the polymer is given by

$$E = \frac{P k_B T \theta^2}{2L} \tag{1}$$

where P is the persistence length; $\theta$ is the angle between the vectors tangent to the curve at two points separated by a distance L; T is the temperature; and $k_B$ the Boltzmann constant. From this expression, the probability of finding a given bending angle is simply

$$\mathcal{P}(\theta) = \sqrt{\frac{P}{2\pi L}} e^{-\frac{P\theta^2}{2L}} \tag{2}$$

From here one can easily obtain the mean cosine of the bending angle, which is given by the following integral

$$< \cos\theta >_{WLC} = \int_{-\infty}^{\infty} d\theta \cos\theta \, \mathcal{P}(\theta) = e^{-L/_{2P}} \tag{3}$$

and then the mean squared end-to-end distance

$$< R^2 > = \int_0^L ds \int_0^L ds' < \cos(\theta(s,s')) > = 2\int_0^L ds \int_0^s ds' e^{\frac{-(s-s')}{2P}} = 4PL\left[1 - \frac{2P}{L}(1 - e^{-L/2P})\right] \tag{4}$$

#### 2. Intrinsically-bent WLC

For an is intrinsically bent molecule, the minimum energy does not correspond to zero bending angle. Thus, we will assume that the intrinsic bending describes a circle of radius $R_0$ and curvature $a \equiv 1/R_0$. In this case, the energy reads

$$E = \frac{P k_B T (\theta - aL)^2}{2L} \tag{5}$$

Then, the angle probability is

$$\mathcal{P}(\theta) = \sqrt{\frac{P}{2\pi L}} e^{-\frac{P(\theta - aL)^2}{2L}} \tag{6}$$

And the mean cosine can be obtained as

$$< \cos\theta > = \sqrt{\frac{P}{2\pi L}} \int_{-\infty}^{\infty} d\theta \cos\theta \, e^{-\frac{P(\theta - aL)^2}{2L}} = \sqrt{\frac{P}{2\pi L}} \int_{-\infty}^{\infty} d\theta \cos(\theta + aL) \, e^{-\frac{P\theta^2}{2L}} = \cos(\mathrm{aL}) \, e^{-\frac{L}{2P}}$$

(7)

This expression is shown in the main text and was used to fit the A-tracts AFM data. The formula for the mean squared end-to-end distance can be derived in the same way as done for the WLC. In this case, one would need to solve the following integral

$$< R^2 > = 2\int_0^L ds \int_0^s ds' \cos(a(s-s'))\, e^{\frac{-(s-s')}{2P}}$$

(8)

This integral is more cumbersome than for the WLC case, but can also be solved analytically yielding the expression shown in the main text

$$< R^2_{s,s+L} >= \frac{2}{a^2+b^2}\left\{bL + \frac{1}{a^2+b^2}\left[(a^2-b^2)(1-cos(aL)e^{-bL}) - 2absin(aL)e^{-bL}\right]\right\}$$

(9)

with $b \equiv 1/(2P_0)$. Notice that by making a=0, one recovers the expression of the WLC, as required.

However, there is a more elegant solution to the problem. Realize that

$$< \cos\theta > = \mathcal{R}e\left\{e^{-\frac{L}{2P^*}}\right\}$$

(10)

Where $\mathcal{R}e$ denotes the real part and $P^*$ is a complex number given by

$$P^* = \frac{P}{1-2iPa}$$

(11)

The mean squared end-to-end distance is then obtained in a straightforward way, following the same rationale employed for the integrals of the WLC

$$< R^2 >= \int_0^L ds \int_0^L ds' \mathcal{R}e\left\{e^{-\frac{|s-s'|}{2P^*}}\right\} = \mathcal{R}e\left\{\int_0^L ds \int_0^L ds'\, e^{-\frac{|s-s'|}{2P^*}}\right\} = \mathcal{R}e\left\{4P^*L\left[1-\frac{2P^*}{L}(1-e^{-\frac{L}{2P^*}})\right]\right\}$$

(12)

This expression reveals that the mean squared end-to-end distance in the IBWLC is simply the real part of the one from the WLC when a complex persistence length is used. We will express this as

$$< R^2_{IBWLC} >= Re < R^2_{WLC;P\to P^*} >$$

(13)

Expanding Eq. (12) one arrives at Eq. (9).

#### Determining the role of monovalent ions on A-tract bending

As an indirect readout of the global curvature of A-tracts, we obtained force-extension curves at different ionic conditions, from where we determined $\mathrm{P}$ (Fig. 3b, Table 1). We found a reduction of the persistence length at higher concentrations of sodium ions, a trend already described for random sequence DNA. The persistence length can be decomposed in two terms as $\mathrm{P} = \mathrm{P}_{\infty} + \mathrm{P}_{\mathrm{el}}(\mathrm{c})$, being $\mathrm{P}_{\infty}$ the intrinsic persistence length, and $\mathrm{P}_{\mathrm{el}}(\mathrm{c})$ a electrostatic persistence length dependent on the ionic concentration, $\mathrm{c}$ [3]. The electrostatic component is given by $\mathrm{P}_{\mathrm{el}}(\mathrm{c}) = \mathrm{mc}^{\upsilon}$, where the exponent can take values of $\upsilon$ = -1 [4], or or $\upsilon$ = -1/2 [5]. It has been shown that this model captures the ionic dependence of the persistence length for dsDNA and dsRNA [6], as well as for ssDNA [7]. We fitted our results for $\mathrm{P}$ by using $\upsilon$ = -1/2, and yielding $P_{\infty}$ = 42 ± 1 nm for control molecules and $P_{\infty}$ = 14 ± 1 nm for A tracts. These values set the asymptotic limit for high salt concentration, indicating that Na ions do not reduce significantly the curvature of A-tracts. In addition, the trend for $\mathrm{P}$ obtained for A tracts in the full range of salt concentrations is similar to that of control DNAs ($\mathrm{m}_{\mathrm{A-tract}} = 11 \pm 4$ nm·mM$^{1/2}$ and $\mathrm{m}_{\mathrm{Control}} = 7 \pm 2$ nm·mM$^{1/2}$). This reflects the change in bendability associated to the screening of charges of the phosphate backbone and indicates that A-tracts are intrinsically bent in an ion-independent manner.

#### Supporting Figures

**Figure S1. Migration of control and A-tracts substrates in agarose gels.** 0.8 % agarose gel electrophoresis of AFM substrates (control DNA (2645 bp) lane 2 and A-tracts (2636 bp) lane 3), and central insert before ligation of the tailed oligos of magnetic and optical tweezers substrates (control, lane 4 and A-tracts, lane 5 (5272 bp)) were run 30 ng/lane). Lanes 1 and 6 correspond to 1 kbp DNA ladder. The electrophoresis was run at room temperature for 50 min at 90 V in the absence of intercalator and the gel was later stained with SYBR®Safe. The substrates containing the A-tracts migrated slower than one would expect on the basis of its length.

**Figure S2. Magnetic Tweezers force-extension curves of A-tracts and control molecules at different NaCl concentrations.** Average force-extension curves of A-tracts and control molecules obtained at different concentrations of NaCl. Error bars are standard error of the mean. The data were fitted to the WLC formula (Eq. 7, main text). The values of the fitting parameters are represented in **Fig. 3b, c** main text.

**Figure S3. Overstretching transition of control and A-tracts molecules.** Representative force-extension curves showing the overstretching transitions of control (red) and A-tracts (blue) at forces ~60 pN.

**Figure S4. Representative optical tweezers force-extension curves of A-tracts and control molecules at different NaCl concentrations.** All data sets were fitted to the extensible WLC (eWLC) in the 10-45 pN force range. The data and the fit at 100 mM NaCl is the same as in **Fig. 4a**, main text. The values of the fits shown at 50 mM NaCl were $L_0 = 1834$ nm, $P = 45$ nm, $S = 1440$ pN and $L_0 = 1828$ nm, $P = 48$ nm, $S = 2545$ pN for the control and A-tracts, respectively; and the ones at 500 mM NaCl were $L_0 = 1841$ nm, $P = 34$ nm, $S = 1675$ pN and $L_0 = 1833$ nm, $P = 35$ nm, $S = 2286$ pN for the control and for the A-tracts.

#### Supporting Tables

**Supplementary Table S1.** *DNA primers used in this work.*

| Fragment | Oligonucleotide | Sequence |
|---|---|---|
| Intron | 58.F Bam-Xho-Psi intron4 | GCGTAAGTGGATCCCTCGAGTTATAACAGGTAGCGGAGAAATTG AAG |
| | 59.RApa-Eco-Sal intron4 | ACTTACGCGGGCCCGATATCGTCGACGGTCCCTGTAAGAAATTT TAAAGG |
| Control tweezers substrate | 89.F lambda 40002 XhoI | GCGTAAGTCTCGAGCCGGATGCGGAGTCTTATCC |
| | 90.R lambda 45263 ApaI | GCGTAAGTGGGCCCCGCAAGGATTGCCCCGATG |
| Digoxigenin tailed oligos | XbaI-A | [pho]CTAGACCCGGGCTCGAGGATCCCC |
| | 88.XbaI C ApaI | GGGGATCCTCGAGCCCGGGTCTAGGGCC |
| Biotin tailed oligos | 27P-XhoI-A | [pho]TCGAGCCCGGGCCATGGGATCCCC |
| | 26XhoI-B | GGGGATCCCATGGCCCGGGC |

**Supplementary Table S2.** *Sequence of DNA fragments used in this work.* Groups or two or more consecutive adenines were highlighted in green and groups of two or more thymines are shown in red. All sequences are written from the 5' end.

| Fragment | Size (bp) | Sequence |
|---|---|---|
| A-tract AFM substrate | 2636 | TCGAGTTATAACAGGTAGCGGAGAAATTGAAGAAAAATGGTCAAAAAATCGATGAAAAAC<br>CCGAAAAACTCACGAAAATTCATTAAAATCTTAAAAATTGCTAGAAATTCGGAAAAATGC<br>CCAAAATTTTAGAAAATGTTTAAATATTCCTGAGAAAATGTCAAAAATACCGAAATTTAT<br>TAGTAAAATCTTGAAAATTTTGCCAAAAATCTGGAAAAACCTGGAATTTTCGACTTTTTT<br>TTTCTAAAAGTTTCAAAATTCCCAAATAATCATTCAAAAAAAAAAACTTTCTAAAAAATC<br>TCTAAAATTTTTGCGGAAATTTTGAAAAAATTCACTAAAAAATTGAAATATCCCCGGAAA<br>TTTCGAGAAAATCCAAAAATTTCGGAAAATACACAAAAATACCGGTTTTTGCTGAAAAAG<br>CACAGAAGTTTTGCGACAAAACCTTTAAAAATCTGTAAAATCCCCCAAAAATTTTCAAAT<br>TTTTTCTCGAAAAATTGCAAAAAATGTAAAAATTCCCCCAAATAATCCGAAAATCCAAAA<br>AAATTACCTAAAAATCGGGAAAATTTTGAAAAATCCCTGAAAATTGTCGTAAAAATACCA<br>AAAATTTCGCTAAAAATCTGAAAAATTGCTAGAAATTCGAAAAAAAATGCTTAAAATTTC<br>AGAAAATGTTAAAAATTCCCAAAAAACATGTAAAATCTTGAGAAAATGTCAAAAATACCG<br>AAATTTATTAGAAAAATCTGAAAACTCACTGACATTTTGCGAAAAATTCCGAAAATTGGT<br>TAAAAAATTTTTTTAAAAAACCTGGAATTTTCCAAATTTTTTTTTCTAAAAATTAAAAAA<br>TTCCGAAAAATCCTTTAAAATTTCTTACAGGGACCGTCGAGTTATAACAGGTAGCGGAGA<br>AATTGAAGAAAAATGGTCAAAAAATCGATGAAAAACCCGAAAAACTCACGAAAATTCATT<br>AAAATCTTAAAAATTGCTAGAAATTCGGAAAAATGCCCAAAATTTTAGAAAATGTTTAAA<br>TATTCCTGAGAAAATGTCAAAAATACCGAAATTTATTAGTAAAATCTTGAAAATTTTGCC<br>AAAAATCTGGAAAAACCTGGAATTTTCGACTTTTTTTTTCTAAAAGTTTCAAAATTCCCA<br>AATAATCATTCAAAAAAAAAAACTTTCTAAAAAATCTCTAAAATTTTTGCGGAAATTTTG<br>AAAAAATTCACTAAAAAATTGAAATATCCCCGGAAATTTCGAGAAAATCCAAAAATTTCG<br>GAAAATACACAAAAATACCGGTTTTTGCTGAAAAAGCACAGAAGTTTTGCGACAAAACCT<br>TTAAAAATCTGTAAAATCCCCCAAAAATTTTCAAATTTTTTCTCGAAAAATTGCAAAAAA<br>TGTAAAAATTCCCCCAAATAATCCGAAAATCCAAAAAAAATTACCTAAAAATCGGGAAAAT<br>TTTGAAAAATCCCTGAAAATTGTCGTAAAAATACCAAAAATTTCGCTAAAAATCTGAAAA<br>ATTGCTAGAAATTCGAAAAAAAATGCTTAAAATTTCAGAAAATGTTAAAAATTCCCAAAA<br>AACATGTAAAATCTTGAGAAAATGTCAAAAATACCGAAATTTATTAGAAAAATCTGAAAA<br>CTCACTGACATTTTGCGAAAAATTCCGAAAATTGGTTAAAAAATTTTTTTAAAAAACCTG<br>GAATTTTCCAAATTTTTTTTTCTAAAAATTAAAAAATTCCGAAAAATCCTTTAAAATTTC<br>TTACAGGGACCGTCGAGTTATAACAGGTAGCGGAGAAATTGAAGAAAAATGGTCAAAAAA<br>TCGATGAAAAACCCGAAAAACTCACGAAAATTCATTAAAATCTTAAAAATTGCTAGAAAT<br>TCGGAAAAATGCCCAAAATTTTAGAAAATGTTTAAATATTCCTGAGAAAATGTCAAAAAT<br>ACCGAAATTTATTAGTAAAATCTTGAAAATTTTGCCAAAAATCTGGAAAAACCTGGAATT<br>TTCGACTTTTTTTTTCTAAAAGTTTCAAAATTCCCAAATAATCATTCAAAAAAAAAAACT<br>TTCTAAAAAATCTCTAAAATTTTTGCGGAAATTTTGAAAAAATTCACTAAAAAATTGAAA<br>TATCCCCGGAAATTTCGAGAAAATCCAAAAATTTCGGAAAATACACAAAAATACCGGTTT<br>TTGCTGAAAAAGCACAGAAGTTTTGCGACAAAACCTTTAAAAATCTGTAAAATCCCCCAA<br>AAATTTTCAAATTTTTTCTCGAAAAATTGCAAAAAATGTAAAAATTCCCCCAAATAATCC<br>GAAAATCCAAAAAAAATTACCTAAAAATCGGGAAAATTTTGAAAAATCCCTGAAAATTGTC<br>GTAAAAATACCAAAAATTTCGCTAAAAATCTGAAAAATTGCTAGAAATTCGAAAAAAAAT<br>GCTTAAAATTTCAGAAAATGTTAAAAATTCCCAAAAAACATGTAAAATCTTGAGAAAATG |

| | | |
|---|---|---|
| | | TCAAAAATACCGAAATTTATTAGAAAAATCTGAAAACTCACTGACATTTTGCGAAAAATT<br>CCGAAAATTGGTTAAAAAATTTTTTTAAAAAACCTGGAATTTTCCAAATTTTTTTTTCTA<br>AAAATTAAAAAATTCCGAAAAATCCTTTAAAATTTCTTACAGGGACCGTCGACGAT |
| Control AFM substrate | 2645 | TCGACCCTGAAGATGATGTGCTGATGCAGAAAGCGGCAGGGCTTGCCGGAGGTGTCCGCT<br>TTGGCCCGGACGGGAATGAAGTTATCCCCGCTTCCCCGGATGTGGCGGACATGACGGAGG<br>ATGACGTAATGCTGATGACAGTATCAGAAGGGATCGCAGGAGGAGTCCGGTATGGCTGAA<br>CCGGTAGGCGATCTGGTCGTTGATTTGAGTCTGGATGCGGCCAGATTTGACGAGCAGATG<br>GCCAGAGTCAGGCGTCATTTTTCTGGTACGGAAAGTGATGCGAAAAAAACAGCGGCAGTC<br>GTTGAACAGTCGCTGAGCCGACAGGCGCTGGCTGCACAGAAAGCGGGGATTTCCGTCGGG<br>CAGTATAAAGCCGCCATGCGTATGCTGCCTGCACAGTTCACCGACGTGGCCACGCAGCTT<br>GCAGGCGGGCAAAGTCCGTGGCTGATCCTGCTGCAACAGGGGGGGCAGGTGAAGGACTCC<br>TTCGGCGGGATGATCCCCATGTTCAGGGGGCTTGCCGGTGCGATCACCCTGCCGATGGTG<br>GGGGCCACCTCGCTGGCGGTGGCGACCGGTGCGCTGGCGTATGCCTGGTATCAGGGCAAC<br>TCAACCCTGTCCGATTTCAACAAAACGCTGGTCCTTTCCGGCAATCAGGCGGGACTGACG<br>GCAGATCGTATGCTGGTCCTGTCCAGAGCCGGGCAGGCGGCAGGGCTGACGTTTAACCAG<br>ACCAGCGAGTCACTCAGCGCACTGGTTAAGGCGGGGGTAAGCGGTGAGGCTCAGATTGCG<br>TCCATCAGCCAGAGTGTGGCGCGTTTCTCCTCTGCATCCGGCGTGGAGGTGGACAAGGTC<br>GCTGAAGCCTCTAGAGAATGCTACGTACCTGATGAGCTCCAGCTTTTGTTCCCTTTAGTG<br>AGGGTTAATTGCGCGCTTGGCGTAATCATGGTCATAGCTGTTTCCTGTGTGAAATTGTTA<br>TCCGCTCACAATTCCACACAACATACGAGCCGGAAGCATAAAGTGTAAAGCCTGGGGTGC<br>CTAATGAGTGAGCTAACTCACATTAATTGCGTTGCGCTCACTGCCCGCTTTCCAGTCGGG<br>AAACCTGTCGTGCCAGCTGCATTAATGAATCGGCCAACGCGCGGGGAGAGGCGGTTTGCG<br>TATTGGGCGCTCGAGCGCTTCCTCGCTCACTGACTCGCTGCGCTCGGTCGTTCGGCTGCG<br>GCGAGCGGTATCAGCTCACTCAAAGGCGGTAATACGGTTATCCACAGAATCAGGGGATAA<br>CGCAGGAAAGAACATGTGAGCAAAAGGCCAGCAAAAGGCCAGGAACCGTAAAAAGGCCGC<br>GTTGCTGGCGTTTTTCCATAGGCTCCGCCCCCCTGACGAGCATCACAAAAATCGACGCTC<br>AAGTCAGAGGTGGCGAAACCCGACAGGACTATAAAGATACCAGGCGTTTCCCCCTGGAAG<br>CTCCCTCGTGCGCTCTCCTGTTCCGACCCTGCCGCTTACCGGATACCTGTCCGCCTTTCT<br>CCCTTCGGGAAGCGTGGCGCTTTCTCATAGCTCACGCTGTAGGTATCTCAGTTCGGTGTA<br>GGTCGTTCGCTCCAAGCTGGGCTGTGTGCACGAACCCCCCGTTCAGCCCGACCGCTGCGC<br>CTTATCCGGTAACTATCGTCTTGAGTCCAACCCGGTAAGACACGACTTATCGCCACTGGC<br>AGCAGCCACTGGTAACAGGATTAGCAGAGCGAGGTATGTAGGCGGTGCTACAGAGTTCTT<br>GAAGTGGTGGCCTAACTACGGCTACACTAGAAGGACAGTATTTGGTATCTGCGCTCTGCT<br>GAAGCCAGTTACCTTCGGAAAAAGAGTTGGTAGCTCTTGATCCGGCAAACAAACCACCGC<br>TGGTAGCGGTGGTTTTTTTGTTTGCAAGCAGCAGATTACGCGCAGAAAAAAAGGATCTCA<br>AGAAGATCCTTTGATCTTTTCTACGGGGTCTGACGCTCAGTGGAACGAAAACTCACGTTA<br>AGGGATTTTGGTCATGAGATTATCAAAAAGGATCTTCACCTAGATCCTTTTAAATTAAAA<br>ATGAAGTTTTAAATCAATCTAAAGTATATATGAGTAAACTTGGTCTGACAGTTACCAATG<br>CTTAATCAGTGAGGCACCTATCTCAGCGATCTGTCTATTTCGTTCATCCATAGTTGCCTG<br>ACTCCCCGTCGTGTAGATAACTACGATACGGGAGGGCTTACCATCTGGCCCCAGTGCTGC<br>AATGATACCGCGAGACCCACGCTCACCGGCTCCAGATTTATCAGCAATAAACCAGCCAGC<br>CGGAAGGGCCGAGCGCAGAAGTGGTCCTGCAACTTTATCCGCCTCCATCCAGTCTATTAA<br>TTGTTGCCGGGAAGCTAGAGTAAGTAGTTCGCCAGTTAATAGTTTGCGCAACGTTGTTGC<br>CATTGCTACAGGCATCGTGGTGTCACGCTCGTCGTTTGGTATGGCTTCATTCAGCTCCGG<br>TTCCCAACGATCAAGGCGAGTTACATGATCCCCCATGTTGTGCAAAAAAGCGGTTAGCTC<br>CTTCGGTCCTCCGATCGTTGTCAGAAGTAAGTTGGCCGCAGTGTTATCACTCATGGTTAT<br>GGCAGCACTGCATAATTCTCTTACTGTCATGCCATCCGTAAGATGCTTTTCTGTGACTGG<br>TGAGT |
| A-tract tweezers substrate | 5316 | GGGGATCCCATGGCCCGGGCTCGAGTTATAACAGGTAGCGGAGAAATTGAAGAAAAATGG<br>TCAAAAAATCGATGAAAAACCCGAAAAACTCACGAAAATTCATTAAAATCTTAAAAATTG<br>CTAGAAATTCGGAAAAATGCCCAAAATTTTAGAAAATGTTTAAATATTCCTGAGAAAATG<br>TCAAAAATACCGAAATTTATTAGTAAAATCTTGAAAATTTTGCCAAAAATCTGGAAAAAC<br>CTGGAATTTTCGACTTTTTTTTTCTAAAAGTTTCAAAATTCCCAAATAATCATTCAAAAA<br>AAAAAACTTTCTAAAAAATCTCTAAAATTTTTGCGGAAATTTTGAAAAAATTCACTAAAA<br>AATTGAAATATCCCCGGAAATTTCGAGAAAATCCAAAAATTTCGGAAAATACACAAAAAT<br>ACCGGTTTTTGCTGAAAAAGCACAGAAGTTTTGCGACAAAACCTTTAAAAATCTGTAAAA<br>TCCCCCAAAAATTTTCAAATTTTTTCTCGAAAAATTGCAAAAAATGTAAAAATTCCCCCA<br>AATAATCCGAAAATCCAAAAAAATTACCTAAAAATCGGGAAAATTTTGAAAAATCCCTGA<br>AAATTGTCGTAAAAATACCAAAAATTTCGCTAAAAATCTGAAAAATTGCTAGAAATTCGA<br>AAAAAAATGCTTAAAATTTCAGAAAATGTTAAAAATTCCCAAAAAACATGTAAAATCTTG<br>AGAAAATGTCAAAAATACCGAAATTTATTAGAAAAATCTGAAAACTCACTGACATTTTGC<br>GAAAAATTCCGAAAATTGGTTAAAAAATTTTTTTAAAAAACCTGGAATTTTCCAAATTTT<br>TTTTTCTAAAAATTAAAAAATTCCGAAAAATCCTTTAAAATTTCTTACAGGGACCGTCGA<br>GTTATAACAGGTAGCGGAGAAATTGAAGAAAAATGGTCAAAAAATCGATGAAAAACCCGA<br>AAAACTCACGAAAATTCATTAAAATCTTAAAAATTGCTAGAAATTCGGAAAAATGCCCAA<br>AATTTTAGAAAATGTTTAAATATTCCTGAGAAAATGTCAAAAATACCGAAATTTATTAGT<br>AAAATCTTGAAAATTTTGCCAAAAATCTGGAAAAACCTGGAATTTTCGACTTTTTTTTTC<br>TAAAAGTTTCAAAATTCCCAAATAATCATTCAAAAAAAAAAACTTTCTAAAAAATCTCTA<br>AAATTTTTGCGGAAATTTTGAAAAAATTCACTAAAAAATTGAAATATCCCCGGAAATTTC<br>GAGAAAATCCAAAAATTTCGGAAAATACACAAAAATACCGGTTTTTGCTGAAAAAGCACA<br>GAAGTTTTGCGACAAAACCTTTAAAAATCTGTAAAATCCCCCAAAAATTTTCAAATTTTT<br>TCTCGAAAAATTGCAAAAAATGTAAAAATTCCCCCAAATAATCCGAAAATCCAAAAAAAT<br>TACCTAAAAATCGGGAAAATTTTGAAAAATCCCTGAAAATTGTCGTAAAAATACCAAAAA<br>TTTCGCTAAAAATCTGAAAAATTGCTAGAAATTCGAAAAAAAATGCTTAAAATTTCAGAA |

| | | |
|---|---|---|
| | | AATGTTAAAAATTCCCAAAAAACATGTAAAATCTTGAGAAAATGTCAAAAATACCGAAAT<br>TTATTAGAAAAATCTGAAAACTCACTGACATTTTGCGAAAAATTCCGAAAATTGGTTAAA<br>AAATTTTTTTAAAAAACCTGGAATTTTCCAAATTTTTTTTTCTAAAAATTAAAAAATTCC<br>GAAAAATCCTTTAAAATTTCTTACAGGGACCGTCGAGTTATAACAGGTAGCGGAGAAATT<br>GAAGAAAAATGGTCAAAAAATCGATGAAAAACCCGAAAAACTCACGAAAATTCATTAAAA<br>TCTTAAAAATTGCTAGAAATTCGGAAAAATGCCCAAAATTTTAGAAAATGTTTAAATATT<br>CCTGAGAAAATGTCAAAAATACCGAAATTTATTAGTAAAATCTTGAAAATTTTGCCAAAA<br>ATCTGGAAAAACCTGGAATTTTCGACTTTTTTTTTCTAAAAGTTTCAAAATTCCCAAATA<br>ATCATTCAAAAAAAAAAACTTTCTAAAAAATCTCTAAAATTTTTGCGGAAATTTTGAAAA<br>AATTCACTAAAAAATTGAAATATCCCCGGAAATTTCGAGAAAATCCAAAAATTTCGGAAA<br>ATACACAAAAATACCGGTTTTTGCTGAAAAAGCACAGAAGTTTTGCGACAAAACCTTTAA<br>AAATCTGTAAAATCCCCCAAAAATTTTCAAATTTTTTCTCGAAAAATTGCAAAAAATGTA<br>AAAATTCCCCCAAATAATCCGAAAATCCAAAAAAATTACCTAAAAATCGGGAAAATTTTG<br>AAAAATCCCTGAAAATTGTCGTAAAAATACCAAAAATTTCGCTAAAAATCTGAAAAATTG<br>CTAGAAATTCGAAAAAAAATGCTTAAAATTTCAGAAAATGTTAAAAATTCCCAAAAAACA<br>TGTAAAATCTTGAGAAAATGTCAAAAATACCGAAATTTATTAGAAAAATCTGAAAACTCA<br>CTGACATTTTGCGAAAAATTCCGAAAATTGGTTAAAAAATTTTTTTAAAAAACCTGGAAT<br>TTTCCAAATTTTTTTTTCTAAAAATTAAAAAATTCCGAAAAATCCTTTAAAATTTCTTAC<br>AGGGACCGTCGAGTTATAACAGGTAGCGGAGAAATTGAAGAAAAATGGTCAAAAAATCGA<br>TGAAAAACCCGAAAAACTCACGAAAATTCATTAAAATCTTAAAAATTGCTAGAAATTCGG<br>AAAAATGCCCAAAATTTTAGAAAATGTTTAAATATTCCTGAGAAAATGTCAAAAATACCG<br>AAATTTATTAGTAAAATCTTGAAAATTTTGCCAAAAATCTGGAAAAACCTGGAATTTTCG<br>ACTTTTTTTTTCTAAAAGTTTCAAAATTCCCAAATAATCATTCAAAAAAAAAAACTTTCT<br>AAAAAATCTCTAAAATTTTTGCGGAAATTTTGAAAAAATTCACTAAAAAATTGAAATATC<br>CCCGGAAATTTCGAGAAAATCCAAAAATTTCGGAAAATACACAAAAATACCGGTTTTTGC<br>TGAAAAAGCACAGAAGTTTTGCGACAAAACCTTTAAAAATCTGTAAAATCCCCCAAAAAT<br>TTTCAAATTTTTTCTCGAAAAATTGCAAAAAATGTAAAAATTCCCCCAAATAATCCGAAA<br>ATCCAAAAAAATTACCTAAAAATCGGGAAAATTTTGAAAAATCCCTGAAAATTGTCGTAA<br>AAATACCAAAAATTTCGCTAAAAATCTGAAAAATTGCTAGAAATTCGAAAAAAAATGCTT<br>AAAATTTCAGAAAATGTTAAAAATTCCCAAAAAACATGTAAAATCTTGAGAAAATGTCAA<br>AAATACCGAAATTTATTAGAAAAATCTGAAAACTCACTGACATTTTGCGAAAAATTCCGA<br>AAATTGGTTAAAAAATTTTTTTAAAAAACCTGGAATTTTCCAAATTTTTTTTTCTAAAAA<br>TTAAAAAATTCCGAAAAATCCTTTAAAATTTCTTACAGGGACCGTCGAGTTATAACAGGT<br>AGCGGAGAAATTGAAGAAAAATGGTCAAAAAATCGATGAAAAACCCGAAAAACTCACGAA<br>AATTCATTAAAATCTTAAAAATTGCTAGAAATTCGGAAAAATGCCCAAAATTTTAGAAAA<br>TGTTTAAATATTCCTGAGAAAATGTCAAAAATACCGAAATTTATTAGTAAAATCTTGAAA<br>ATTTTGCCAAAAATCTGGAAAAACCTGGAATTTTCGACTTTTTTTTTCTAAAAGTTTCAA<br>AATTCCCAAATAATCATTCAAAAAAAAAAACTTTCTAAAAAATCTCTAAAATTTTTGCGG<br>AAATTTTGAAAAAATTCACTAAAAAATTGAAATATCCCCGGAAATTTCGAGAAAATCCAA<br>AAATTTCGGAAAATACACAAAAATACCGGTTTTTGCTGAAAAAGCACAGAAGTTTTGCGA<br>CAAAACCTTTAAAAATCTGTAAAATCCCCCAAAAATTTTCAAATTTTTTCTCGAAAAATT<br>GCAAAAAATGTAAAAATTCCCCCAAATAATCCGAAAATCCAAAAAAATTACCTAAAAATC<br>GGGAAAATTTTGAAAAATCCCTGAAAATTGTCGTAAAAATACCAAAAATTTCGCTAAAAA<br>TCTGAAAAATTGCTAGAAATTCGAAAAAAAATGCTTAAAATTTCAGAAAATGTTAAAAAT<br>TCCCAAAAAACATGTAAAATCTTGAGAAAATGTCAAAAATACCGAAATTTATTAGAAAAA<br>TCTGAAAACTCACTGACATTTTGCGAAAAATTCCGAAAATTGGTTAAAAAATTTTTTTAA<br>AAAACCTGGAATTTTCCAAATTTTTTTTTCTAAAAATTAAAAAATTCCGAAAAATCCTTT<br>AAAATTTCTTACAGGGACCGTCGAGTTATAACAGGTAGCGGAGAAATTGAAGAAAAATGG<br>TCAAAAAATCGATGAAAAACCCGAAAAACTCACGAAAATTCATTAAAATCTTAAAAATTG<br>CTAGAAATTCGGAAAAATGCCCAAAATTTTAGAAAATGTTTAAATATTCCTGAGAAAATG<br>TCAAAAATACCGAAATTTATTAGTAAAATCTTGAAAATTTTGCCAAAAATCTGGAAAAAC<br>CTGGAATTTTCGACTTTTTTTTTCTAAAAGTTTCAAAATTCCCAAATAATCATTCAAAAA<br>AAAAAACTTTCTAAAAAATCTCTAAAATTTTTGCGGAAATTTTGAAAAAATTCACTAAAA<br>AATTGAAATATCCCCGGAAATTTCGAGAAAATCCAAAAATTTCGGAAAATACACAAAAAT<br>ACCGGTTTTTGCTGAAAAAGCACAGAAGTTTTGCGACAAAACCTTTAAAAATCTGTAAAA<br>TCCCCCAAAAATTTTCAAATTTTTTCTCGAAAAATTGCAAAAAATGTAAAAATTCCCCCA<br>AATAATCCGAAAATCCAAAAAAATTACCTAAAAATCGGGAAAATTTTGAAAAATCCCTGA<br>AAATTGTCGTAAAAATACCAAAAATTTCGCTAAAAATCTGAAAAATTGCTAGAAATTCGA<br>AAAAAAATGCTTAAAATTTCAGAAAATGTTAAAAATTCCCAAAAAACATGTAAAATCTTG<br>AGAAAATGTCAAAAATACCGAAATTTATTAGAAAAATCTGAAAACTCACTGACATTTTGC<br>GAAAAATTCCGAAAATTGGTTAAAAAATTTTTTTAAAAAACCTGGAATTTTCCAAATTTT<br>TTTTTCTAAAAATTAAAAAATTCCGAAAAATCCTTTAAAATTTCTTACAGGGACCGTCGA<br>CGATATCGGGCCCTAGACCCGGGCTCGAGGATCCCC |
| Control tweezers substrate | 5316 | GGGGATCCCATGGCCCGGGcTCGAGCCGGATGCGGAGTCTTATCCGTGGAAATCAAACGC<br>GCACTACTGGCTGGTTACCAACCTGTATCAGAACATGCGGGCCAATGCGCTTACTGATGC<br>GGAATTACGCCGTAAGGCCGCAGATGAGCTTGTCCATATGACTGCGAGAATTAACCGTGG<br>TGAGGCGATCCCTGAACCAGTAAAACAACTTCCTGTCATGGGCGGTAGACCTCTAAATCG<br>TGCACAGGCTCTGGCGAAGATCGCAGAAATCAAAGCTAAGTTCGGACTGAAAGGAGCAAG<br>TGTATGACGGGCAAAGAGGCAATTATTCATTACCTGGGGACGCATAATAGCTTCTGTGCG<br>CCGGACGTTGCCGCGCTAACAGGCGCAACAGTAACCAGCATAAATCAGGCCGCGGCTAAA<br>ATGGCACGGGCAGGTCTTCTGGTTATCGAAGGTAAGGTCTGGCGAACGGTGTATTACCGG<br>TTTGCTACCAGGGAAGAACGGGAAGGAAAGATGAGCACGAACCTGGTTTTTAAGGAGTGT<br>CGCCAGAGTGCCGCGATGAAACGGGTATTGGCGGTATATGGAGTTAAAAGATGACCATCT<br>ACATTACTGAGCTAATAACAGGCCTGCTGGTAATCGCAGGCCTTTTTATTTGGGGGAGAG<br>GGAAGTCATGAAAAAACTAACCTTTGAAATTCGATCTCCAGCACATCAGCAAAACGCTAT<br>TCACGCAGTACAGCAAATCCTTCCAGACCCAACCAAACCAATCGTAGTAACCATTCAGGA |

| | | |
|---|---|---|
| | | ACGCAACCGCAGCTTAGACCAAAACAGGAAGCTATGGGCCTGCTTAGGTGACGTCTCTCG<br>TCAGGTTGAATGGCATGGTCGCTGGCTGGATGCAGAAAGCTGGAAGTGTGTGTTTACCGC<br>AGCATTAAAGCAGCAGGATGTTGTTCCTAACCTTGCCGGGAATGGCTTTGTGGTAATAGG<br>CCAGTCAACCAGCAGGATGCGTGTAGGCGAATTTGCGGAGCTATTAGAGCTTATACAGGC<br>ATTCGGTACAGAGCGTGGCGTTAAGTGGTCAGACGAAGCGAGACTGGCTCTGGAGTGGAA<br>AGCGAGATGGGGAGACAGGGCTGCATGATAAATGTCGTTAGTTTCTCCGGTGGCAGGACG<br>TCAGCATATTTGCTCTGGCTAATGGAGCAAAAGCGACGGGCAGGTAAAGACGTGCATTAC<br>GTTTTCATGGATACAGGTTGTGAACATCCAATGACATATCGGTTTGTCAGGGAAGTTGTG<br>AAGTTCTGGGATATACCGCTCACCGTATTGCAGGTTGATATCAACCCGGAGCTTGGACAG<br>CCAAATGGTTATACGGTATGGGAACCAAAGGATATTCAGACGCGAATGCCTGTTCTGAAG<br>CCATTTATCGATATGGTAAAGAAATATGGCACTCCATACGTCGGCGGCGCGTTCTGCACT<br>GACAGATTAAAACTCGTTCCCTTCACCAAATACTGTGATGACCATTTCGGGCGAGGGAAT<br>TACACCACGTGGATTGGCATCAGAGCTGATGAACCGAAGCGGCTAAAGCCAAAGCCTGGA<br>ATCAGATATCTTGCTGAACTGTCAGACTTTGAGAAGGAAGATATCCTCGCATGGTGGAAG<br>CAACAACCATTCGATTTGCAAATACCGGAACATCTCGGTAACTGCATATTCTGCATTAAA<br>AAATCAACGCAAAAAATCGGACTTGCCTGCAAAGATGAGGAGGGATTGCAGCGTGTTTTT<br>AATGAGGTCATCACGGGATCCCATGTGCGTGACGGACATCGGGAAACGCCAAAGGAGATT<br>ATGTACCGAGGAAGAATGTCGCTGGACGGTATCGCGAAAATGTATTCAGAAAATGATTAT<br>CAAGCCCTGTATCAGGACATGGTACGAGCTAAAAGATTCGATACCGGCTCTTGTTCTGAG<br>TCATGCGAAATATTTGGAGGGCAGCTTGATTTCGACTTCGGGAGGGAAGCTGCATGATGC<br>GATGTTATCGGTGCGGTGAATGCAAAGAAGATAACCGCTTCCGACCAAATCAACCTTACT<br>GGAATCGATGGTGTCTCCGGTGTGAAAGAACACCAACAGGGGTGTTACCACTACCGCAGG<br>AAAAGGAGGACGTGTGGCGAGACAGCGACGAAGTATCACCGACATAATCTGCGAAAACTG<br>CAAATACCTTCCAACGAAACGCACCAGAAATAAACCCAAGCCAATCCCAAAAGAATCTGA<br>CGTAAAAACCTTCAACTACACGGCTCACCTGTGGGATATCCGGTGGCTAAGACGTCGTGC<br>GAGGAAAACAAGGTGATTGACCAAAATCGAAGTTACGAACAAGAAAGCGTCGAGCGAGCT<br>TTAACGTGCGCTAACTGCGGTCAGAAGCTGCATGTGCTGGAAGTTCACGTGTGTGAGCAC<br>TGCTGCGCAGAACTGATGAGCGATCCGAATAGCTCGATGCACGAGGAAGAAGATGATGGC<br>TAAACCAGCGCGAAGACGATGTAAAAACGATGAATGCCGGGAATGGTTTCACCCTGCATT<br>CGCTAATCAGTGGTGGTGCTCTCCAGAGTGTGGAACCAAGATAGCACTCGAACGACGAAG<br>TAAAGAACGCGAAAAAGCGGAAAAAGCAGCAGAGAAGAAACGACGACGAGAGGAGCAGAA<br>ACAGAAAGATAAACTTAAGATTCGAAAACTCGCCTTAAAGCCCCGCAGTTACTGGATTAA<br>ACAAGCCCAACAAGCCGTAAACGCCTTCATCAGAGAAAGAGACCGCGACTTACCATGTAT<br>CTCGTGCGGAACGCTCACGTCTGCTCAGTGGGATGCCGGACATTACCGGACAACTGCTGC<br>GGCACCTCAACTCCGATTTAATGAACGCAATATTCACAAGCAATGCGTGGTGTGCAACCA<br>GCACAAAAGCGGAAATCTCGTTCCGTATCGCGTCGAACTGATTAGCCGCATCGGGCAGGA<br>AGCAGTAGACGAAATCGAATCAAACCATAACCGCCATCGCTGGACTATCGAAGAGTGCAA<br>GGCGATCAAGGCAGAGTACCAACAGAAACTCAAAGACCTGCGAAATAGCAGAAGTGAGGC<br>CGCATGACGTTCTCAGTAAAAACCATTCCAGACATGCTCGTTGAAACATACGGAAATCAG<br>ACAGAAGTAGCACGCAGACTGAAATGTAGTCGCGGTACGGTCAGAAAATACGTTGATGAT<br>AAAGACGGGAAAATGCACGCCATCGTCAACGACGTTCTCATGGTTCATCGCGGATGGAGT<br>GAAAGAGATGCGCTATTACGAAAAAATTGATGGCAGCAAATACCGAAATATTTGGGTAGT<br>TGGCGATCTGCACGGATGCTACACGAACCTGATGAACAAACTGGATACGATTGGATTCGA<br>CAACAAAAAAGACCTGCTTATCTCGGTGGGCGATTTGGTTGATCGTGGTGCAGAGAACGT<br>TGAATGCCTGGAATTAATCACATTCCCCTGGTTCAGAGCTGTACGTGGAAACCATGAGCA<br>AATGATGATTGATGGCTTATCAGAGCGTGGAAACGTTAATCACTGGCTGCTTAATGGCGG<br>TGGCTGGTTCTTTAATCTCGATTACGACAAAGAAATTCTGGCTAAAGCTCTTGCCCATAA<br>AGCAGATGAACTTCCGTTAATCATCGAACTGGTGAGCAAAGATAAAAAATATGTTATCTG<br>CCACGCCGATTATCCCTTTGACGAATACGAGTTTGGAAAGCCAGTTGATCATCAGCAGGT<br>AATCTGGAACCGCGAACGAATCAGCAACTCACAAAACGGGATCGTGAAAGAAATCAAAGG<br>CGCGGACACGTTCATCTTTGGTCATACGCCAGCAGTGAAACCACTCAAGTTTGCCAACCA<br>AATGTATATCGATACCGGCGCAGTGTTCTGCGGAAACCTAACATTGATTCAGGTACAGGG<br>AGAAGGCGCATGAGACTCGAAAGCGTAGCTAAATTTCATTCGCCAAAAAGCCCGATGATG<br>AGCGACTCACCACGGGCCACGGCTTCTGACTCTCTTTCCGGTACTGATGTGATGGCTGCT<br>ATGGGGATGGCGCAATCACAAGCCGGATTCGGTATGGCTGCATTCTGCGGTAAGCACGAA<br>CTCAGCCAGAACGACAAACAAAAGGCTATCAACTATCTGATGCAATTTGCACACAAGGTA<br>TCGGGGAAATACCGTGGTGTGGCAAAGCTTGAAGGAAATACTAAGGCAAAGGTACTGCAA<br>GTGCTCGCAACATTCGCTTATGCGGATTATTGCCGTAGTGCCGCGACGCCGGGGGCAAGA<br>TGCAGAGATTGCCATGGTACAGGCCGTGCGGTTGATATTGCCAAAACAGAGCTGTGGGGG<br>AGAGTTGTCGAGAAAGAGTGCGGAAGATGCAAAGGCGTCGGCTATTCAAGGATGCCAGCA<br>AGCGCAGCATATCGCGCTGTGACGATGCTAATCCCAAACCTTACCCAACCCACCTGGTCA<br>CGCACTGTTAAGCCGCTGTATGACGCTCTGGTGGTGCAATGCCACAAAGAAGAGTCAATC<br>GCAGACAACATTTTGAATGCGGTCACACGTTAGCAGCATGATTGCCACGGATGGCAACAT<br>ATTAACGGCATGATATTGACTTATTGAATAAAATTGGGTAAATTTGACTCAACGATGGGT<br>TAATTCGCTCGTTGTGGTAGTGAGATGAAAAGAGGCGGCGCTTACTACCGATTCCGCCTA<br>GTTGGTCACTTCGACGTATCGTCTGGAACTCCAACCATCGCAGGCAGAGAGGTCTGCAAA<br>ATGCAATCCCGAAACAGTTCGCAGGTAATAGTTAGAGCCTGCATAACGGTTTCGGGATTT<br>TTTATATCTGCACAACAGGTAAGAGCATTGAGTCGATAATCGTGAAGAGTCGGCGAGCCT<br>GGTTAGCCAGTGCTCTTTCCGTTGTGCTGAATTAAGCGAATACCGGAAGCAGAACCGGAT<br>CACCAAATGCGTACAGGCGTCATCGCCGCCCAGCAACAGCACAACCCAAACTGAGCCGTA<br>GCCACTGTCTGTCCTGAATTCATTAGTAATAGTTACGCTGCGGCCTTTTACACATGACCT<br>TCGTGAAAGCGGGTGGCAGGAGGTCGCGCTAACAACCTCCTGCCGTTTTGCCCGTGCATA<br>TCGGTCACGAACAAATCTGATTACTAAACACAGTAGCCTGGATTTGTTCTATCAGTAATC<br>GACCTTATTCCTAATTAAATAGAGCAAATCCCCTTATTGGGGGTAAGACATGAAGATGCC<br>AGAAAAACATGACCTGTTGGCCGCCATTCTCGCGGCAAAGGAACAAGGCATCGGGGCAAT<br>CCTTGCGGGGCCCTAGACCCGGGCTCGAGGATCCCC |

#### Figure S1

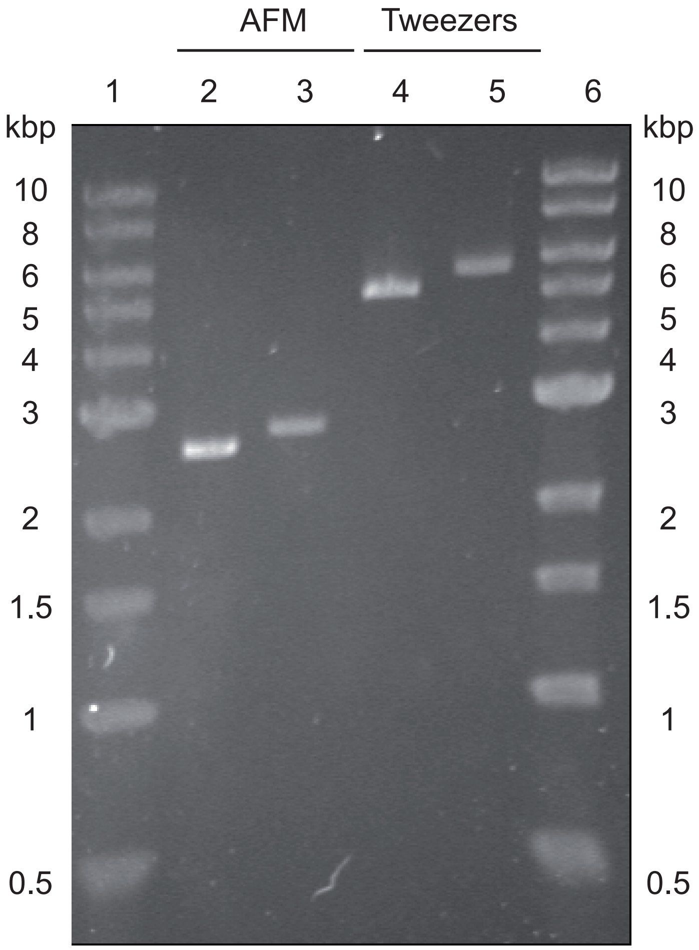

Figure S2

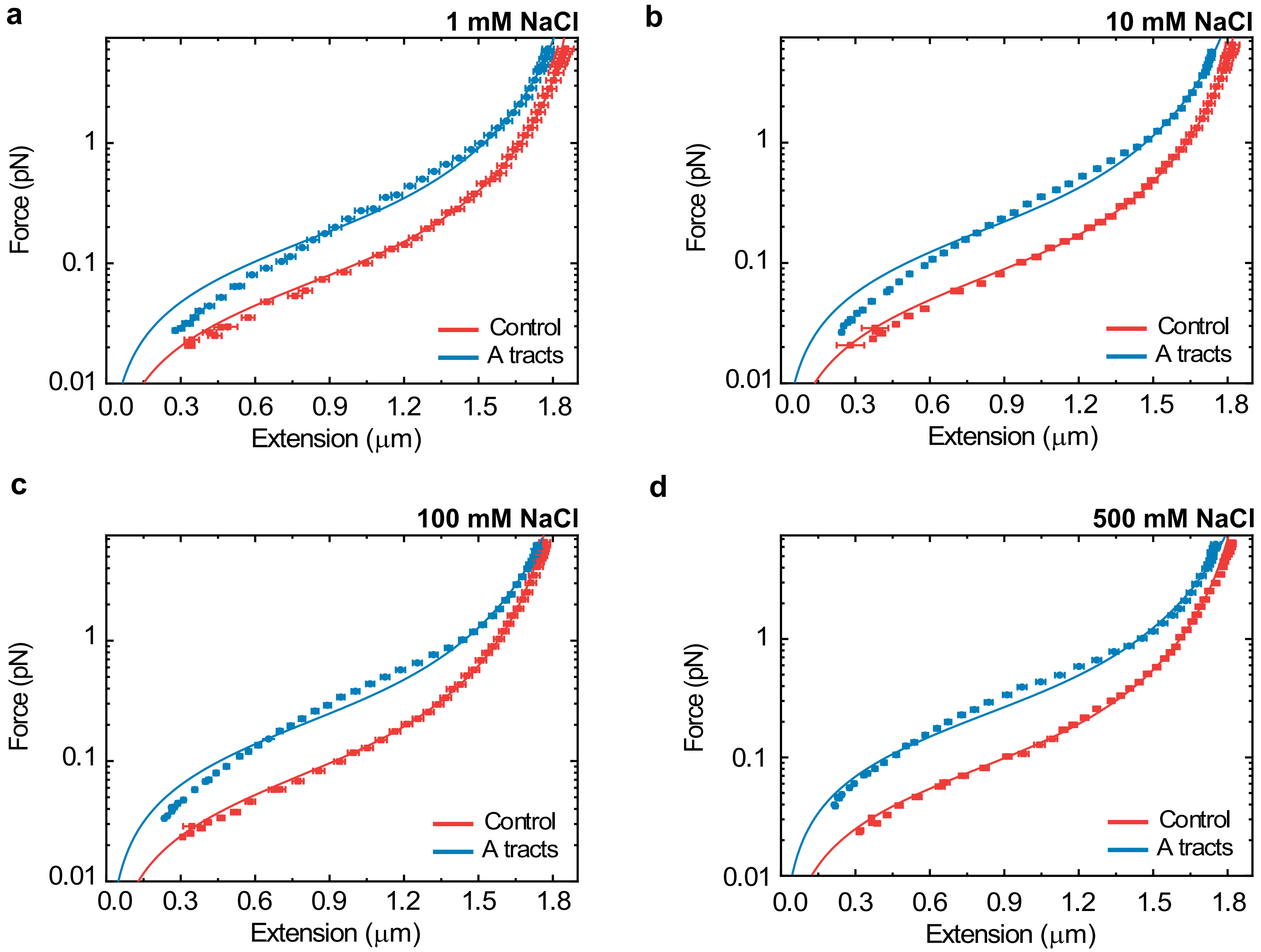

a
1 mM NaCl
b
10 mM NaCl
c
100 mM NaCl
d
500 mM NaCl
Force (pN)
Extension (μm)
Control
A tracts
0.01
0.1
1
0.0
0.3
0.6
0.9
1.2
1.5
1.8

#### Figure S3

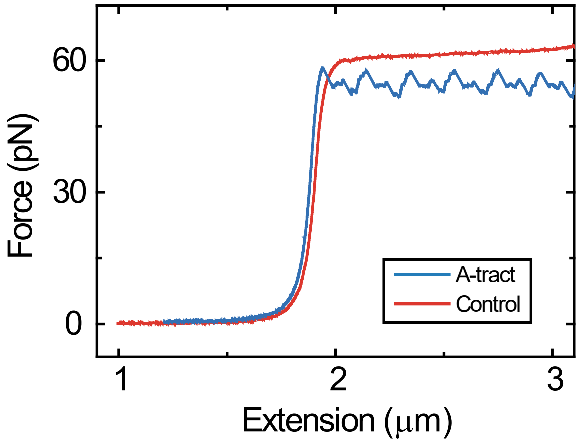

**Figure S4**

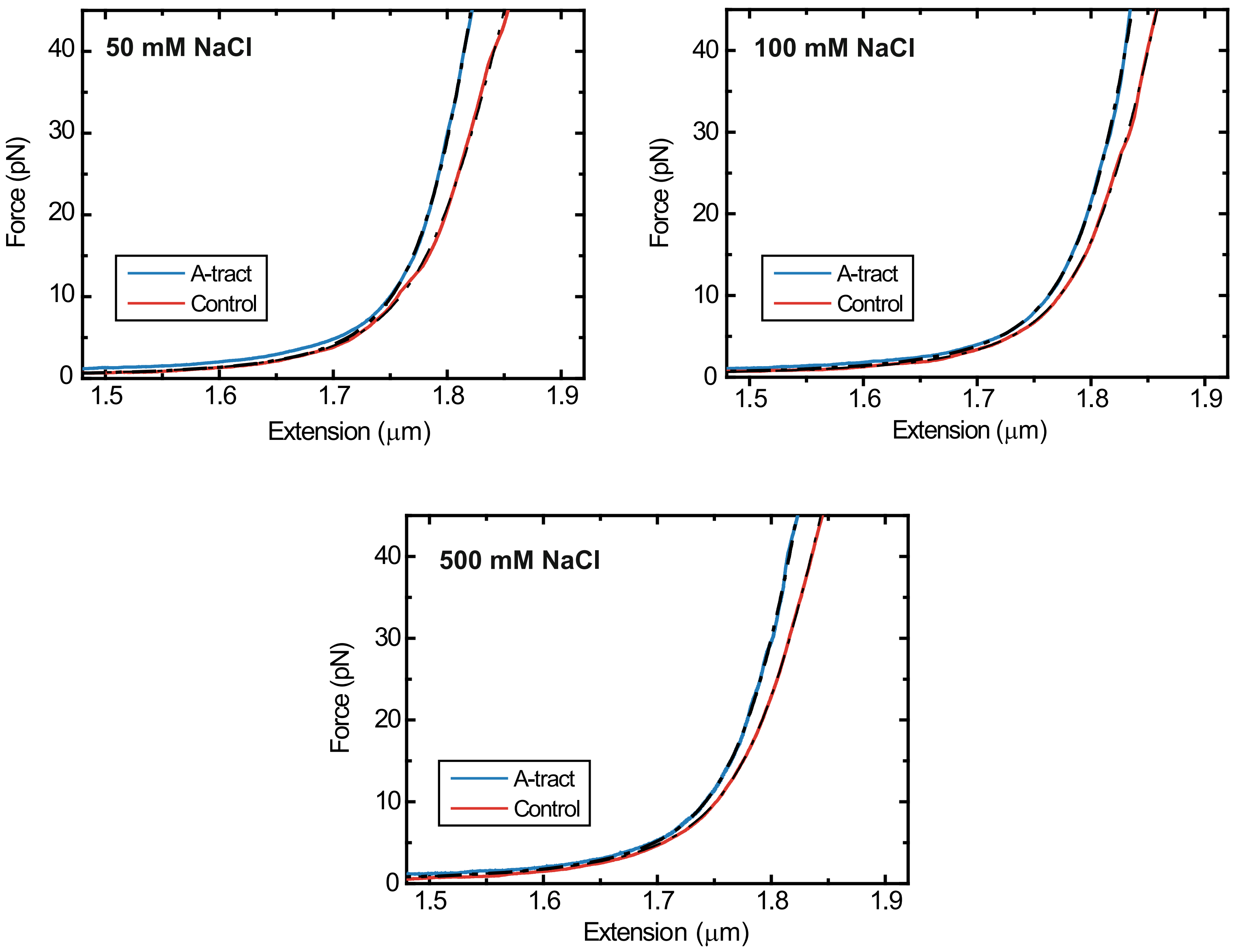

50 mM NaCl
Force (pN)
Extension (μm)
A-tract
Control
100 mM NaCl
Force (pN)
Extension (μm)
A-tract
Control
500 mM NaCl
Force (pN)
Extension (μm)
A-tract
Control
